# Transmission dynamics and pathogenesis differ between pheasants and partridges infected with clade 2.3.4.4b H5N8 and H5N1 high-pathogenicity avian influenza viruses

**DOI:** 10.1101/2023.09.22.558959

**Authors:** Amanda H. Seekings, Yuan Liang, Caroline J. Warren, Charlotte K. Hjulsager, Saumya S. Thomas, Fabian Z. X. Lean, Alejandro Núñez, Paul Skinner, David Selden, Marco Falchieri, Hugh Simmons, Ian H. Brown, Lars. E. Larsen, Ashley C. Banyard, Marek J. Slomka

## Abstract

During the UK 2020-2021 epizootic of H5Nx clade 2.3.4.4b high-pathogenicity avian influenza viruses (HPAIVs), high mortality occurred during incursions in commercially farmed common pheasants (*Phasianus colchicus)*. Two pheasant farms, affected separately by H5N8 and H5N1 subtypes, included adjacently housed red-legged partridges (*Alectoris rufa*), which appeared to be unaffected. Despite extensive ongoing epizootics, H5Nx HPAIV partridge outbreaks were not reported during 2020-2021 and 2021-2022 in the UK, so it is postulated that partridges are more resistant to HPAIV infection than other gamebirds. To assess this, pathogenesis and both intra- and inter-species transmission of UK pheasant-origin H5N8-2021 and H5N1-2021 HPAIVs were investigated. Onward transmission to chickens was also assessed to better understand the risk of spread from gamebirds to other commercial poultry sectors. A lower infectious dose was required to infect pheasants with H5N8-2021 compared to H5N1-2021. However, HPAIV systemic dissemination to multiple organs within pheasants was more rapid following infection with H5N1-2021 than H5N8-2021, with the former attaining generally higher viral RNA levels in tissues. Intraspecies transmission to contact pheasants was successful for both viruses and associated with viral environmental contamination, while interspecies transmission to a first chicken-contact group was also efficient. However, further onward transmission to additional chicken contacts was only achieved with H5N1-2021. Intra-partridge transmission was only successful when high-dose H5N1-2021 was administered, while partridges inoculated with H5N8-2021 failed to shed and transmit, although extensive tissue tropism was observed for both viruses. Mortalities among infected partridges featured a longer incubation period compared to that in pheasants, for both viruses. Therefore, the susceptibility of different gamebird species and pathogenicity outcomes to the ongoing H5Nx clade 2.3.4.4b HPAIVs varies, but pheasants represent a greater likelihood of H5Nx HPAIV introduction into galliforme poultry settings. Consequently, viral maintenance within gamebird populations and risks to poultry species warrant enhanced investigation.

## Introduction

Avian Influenza (AI) is caused by influenza A viruses affecting many wild and domesticated avian species [1]. AI virus (AIV) isolates are classified, based on their external glycoproteins, into 16 hemagglutinin (HA: H1-H16) and nine neuraminidase (NA: N1-N9) subtypes and continue to evolve both through genetic drift [2] and reassortment of their gene segments [3]. Many AIVs persist in wild anseriforme populations as low-pathogenicity avian influenza viruses (LPAIVs), causing mainly subclinical infections [4]. Viral spill-over of H5 and H7 LPAIV subtypes into galliforme hosts such as chickens can result in a switch from LPAIV to high-pathogenicity (HP)AIV, with high mortality and associated significant economic losses [5].

In 1996 a novel H5N1 HPAIV “goose/Guangdong” (GsGd) lineage emerged in China, which has subsequently evolved into different clades classified according to their HA (H5) gene phylogeny [6]. Reassortments which include NA (and internal) gene exchanges have featured in the evolution of clade 2.3.4.4 GsGd HPAIVs which are also referred to as H5Nx HPAIVs [7]. European clade 2.3.4.4 H5Nx HPAIV incursions have occurred since 2014 during the autumn / winter [8] with ensuing seasonal subtype dominance; namely H5N8 during 2014-2015, 2016-2017, 2019-2020 [9,10] and H5N6 during 2018 [11]. Reassortments which include NA (and internal) gene exchanges featured in the evolution of clade 2.3.4.4 GsGd HPAIVs which are also referred to as H5Nx HPAIVs [7]. European clade 2.3.4.4 H5Nx HPAIV incursions have occurred since 2014 during the autumn / winter [8] with ensuing seasonal subtype dominance; namely H5N8 during 2014-2015, 2016-2017, 2019-2020 [9,10], H5N6 during 2018 [11] and H5N1 during the 2021-2022 epizootics [12]. However, multiple subtypes can co-circulate as during the 2020-2021 season when H5N8 predominated, wherea minority H5N1 reassortant featured six viral gene changes, namely the NA and five internal gene segments [12–14]. Following H5Nx GsGd HPAIV infection, wild and domesticated anseriformes display a more ameliorated pathogenesis compared to galliformes, hence ducks serve as an insidious reservoir with migratory waterfowl disseminating new infections [15,16].

In the United Kingdom (UK) and Europe, common pheasants (also known as ring-necked pheasants, *Phasianus colchicus*) and red-legged partridges (*Alectoris rufa*) are gallinaceous species reared for recreational hunting, hence are classified as poultry before being released into the wild [17–19]. Juvenile gamebirds are kept as multi-species, multi-age and high-density populations on premises with limited biosecurity [20], with estimates that between 39-57 million pheasants and 8.1-13 million partridges are released annually in the UK [18,21]. The introduction of such a large biomass into the ecosystem impacts on habitats and wildlife, with accompanying challenges to infectious disease control. This biological overabundance may affect the infection status of wild birds by increasing pathogen circulation. Contact between gamebirds and potential AIV reservoir species such as waterfowl is possible during their juvenile phase, with 500,000-1 million mallard ducks (*Anas platyrhynchos*) also raised for shooting in the UK, with released gamebirds sharing foraging locations with wild birds [22]. Therefore, it is important to investigate the role of gamebirds in AIV circulation and transmission, including a potential bridging role for AIVs between wild birds and commercial poultry [23].

Experimental infection of pheasants with North American H5N8 and H5N2 clade 2.3.4.4 HPAIVs isolated in late 2014 caused up to 100% mortality with successful transmission of both subtypes to in-contact pheasants [24], which contrasted with subclinical infection outcomes with a historic unrelated North American H5 HPAIV [25]. The European H5N8 clade 2.3.4.4b HPAIV epizootic during 2016-2017 included UK pheasant farm outbreaks which featured frequent mortality [20]. More recently, the clade 2.3.4.4b H5N6 HPAIV cases in Danish wild birds during summer 2018 included detections in wild pheasants which again drew attention to their potential bridging role, although no commercial chicken premises were affected [26]. In an *in vivo* investigation, H5N6 HPAIV direct infection of pheasants initiated efficient onward transmission via introduced contact pheasants, despite high mortality [27]. H5Nx HPAIV caused high mortalities in farmed pheasants in the UK, Denmark and Finland during the 2020-2021 and 2021-2022 clade 2.3.4.4b epizootics, but very few infected wild partridges were reported [28–32]. Direct experimental inoculation of closely related chukar partridges (*Alectoris chukar*) with both the North American (2014) H5N8 and H5N2 clade 2.3.4.4 HPAIVs also caused mortality [24].

Differing species-specific outcomes in gamebirds were inferred at H5Nx clade 2.3.4.4b HPAIV outbreaks during early 2021, with two mixed gamebird premises affected by the H5N8 and H5N1 subtypes in Wales and Scotland, respectively [28]. H5Nx viral shedding with accompanying mortality was only observed among pheasants at both farms, but not for the adjacently housed partridges which remained apparently healthy at statutory culling (APHA, unpublished observations). The classification of farmed gamebirds as poultry necessitates monitoring of H5 and H7 AIV incursions into this sector, where statutory interventions are required to control notifiable avian disease [33]. Therefore, we investigated experimental infections of pheasants and partridges with the UK pheasant-origin H5N8 and H5N1 HPAIVs. Transmission outcomes for both subtypes were compared in both gamebird species, along with potential onward spread to chickens. Pathogenesis and transmission dynamics were assessed to understand species-specific differences between gamebirds in their possible role as bridging hosts for contemporary clade 2.3.4.4b viruses.

## Materials and Methods

### Animals: Ethical and safety considerations

Gamebirds (the common pheasant (*P. colchicus*) and red-legged partridge (*A. rufa*)) and specified-pathogen free (SPF) white leghorn chickens (*Gallus domesticus)* were sourced from Hy-Fly Game Hatcheries (UK) and Valo (Hungary), respectively. Swabs (buccal and cloacal) and blood were collected prior to infection or exposure to the viruses to demonstrate freedom from prior or ongoing AIV infection by real-time reverse transcriptase PCR (RRT-PCR) and serological testing. Bird groups were housed in pens (200×100×216cm) which included access to water (replaced daily) and food, with environmental enrichment.

All *in vivo* experiments and procedures were approved by the Animal and Plant Health Agency (APHA-Weybridge) Animal Welfare and Ethical Review Body to ensure compliance with European and UK legislation under the Animal (Scientific Procedures) Act 1986 (ASPA) [34], adhering to UK Home Office project license PP7633638. Welfare inspections were carried out up to three times daily to identify humane endpoints to minimize numbers of “found dead” cases [35], with birds that developed severe clinical disease euthanized immediately by cervical dislocation. Apparently healthy birds which survived to study end were similarly culled. Any culls necessitated by welfare considerations which were unrelated to viral infections (e.g. leg injuries, or where one bird in a given study group had survived and should no longer remain in isolation) were excluded from the Kaplan-Meier survival graphs and associated mean death time (MDT) calculations. All work with potentially H5Nx HPAIV infectious materials and infected birds was carried out in licensed BSL3 laboratory and animal housing facilities at APHA-Weybridge.

### Viruses

The H5N8 clade 2.3.4.4b HPAIV, A/pheasant/Wales/9383/2021 (herewith referred to as H5N8-2021), was isolated from pooled brain tissue from five pheasants from a gamebird breeding farm outbreak in January 2021 [28]. Isolation was carried out by standard methods using 9-10-day-old SPF embryonated chicken eggs (ECEs, Valo), producing first (P1) and then second passage (P2) stocks [36]. The H5N1 clade 2.3.4.4b HPAIV, A/pheasant/Scotland/11039/2021 (herewith referred to as H5N1-2021), was similarly isolated and propagated from pooled mixed viscera (heart, liver, spleen and kidney) from five pheasants from a mixed gamebird breeding premises outbreak in February 2021 [28]. The P2 stocks were filtered (0.2 µm filter; Whatman, Gottingen, Germany) and titrated in ECEs to determine the 50% egg infectious dose (EID_50_) [37], and diluted accordingly for the *in vivo* inoculations in sterile 0.1M pH 7.2 phosphate buffered saline.

### Experimental design

Studies were undertaken with birds at six-eight weeks age to (I) determine the 50% minimum infectious dose (MID_50_) of both H5N8-2021 and H5N1-2021 in pheasants and partridges; (II) investigate pathogenesis of each virus following pheasant or partridge infection; and (III) examine the transmission dynamics of H5N8-2021 and H5N1-2021 from infected pheasants to contact chickens, or from infected pheasants to contact partridges which (if successful) could be extended for onward transmission to chickens.

#### (I) Minimum infectious dose (MID_50_)

To determine the MID_50_ of H5N8-2021 and H5N1-2021 in a given gamebird species, six groups of six birds were divided into separate pens. Three different doses of each virus were administered via the ocular-nasal route, namely 100 µl of low (1×10^2^ EID_50_), medium (1×10^4^ EID_50_) and high (1×10^6^ EID_50_) doses per bird. The H5N6 clade 2.3.4.4 HPAIV study [27] suggested that pheasant-to-pheasant transmission of H5N8-2021 and H5N1-2021 was likely to succeed to at least some degree, but attempted transmission among red-legged partridges represented some uncertainty. Therefore, the MID_50_ investigation also provided an opportunity to introduce six naïve contact (recipient, R1) partridges for cohousing at 1-day post-infection (dpi) with the six high-dose directly-inoculated (donor, D0) partridges. Daily swabs (buccal and cloacal) were collected during the MID_50_ experiments.

#### (II) Pathogenesis

Virus dissemination during H5N8-2021 and H5N1-2021 infections were investigated during by time-course experiments. Groups of six pheasants and six partridges were infected with high doses of H5N8-2021 or H5N1-2021. Two birds from each group were planned to be sacrificed at 1, 2, and 3 dpi with tissues collected for viral investigation. Tissue sampling was also performed for any opportunistic mortalities (found dead and euthanised birds) during (I) and (III).

#### (III) Transmission

To investigate the transmission dynamics of both HPAIVs among gamebirds and from gamebirds to chickens, four transmission chains were attempted in four experimental units (Figure 1). All transmission chains were initiated with donor (D0) pheasants being directly inoculated via the ocular-nasal route with 100 µl of high-dose H5N8-2021 or H5N1-2021. Viral shedding was assessed daily by viral RNA (vRNA) testing of buccal and cloacal swabs. At 1 dpi, the first contact group of age-matched recipient birds (R1) were introduced and co-housed with D0 pheasants. At the first detection of vRNA positive swabs from R1 birds, the D0 birds were removed and replaced by naïve (R2) birds, and so forth (Figure 1). H5N8-2021 and H5N1-2021 transmission within pheasants and onwards to chickens was investigated in Transmission Chains A.1 and A.2, respectively. H5N8-2021 and H5N1-2021 transmission from pheasants to partridges and onwards to partridges was examined in Transmission Chains B.1 and B.2, respectively.

**Fig. 1.**
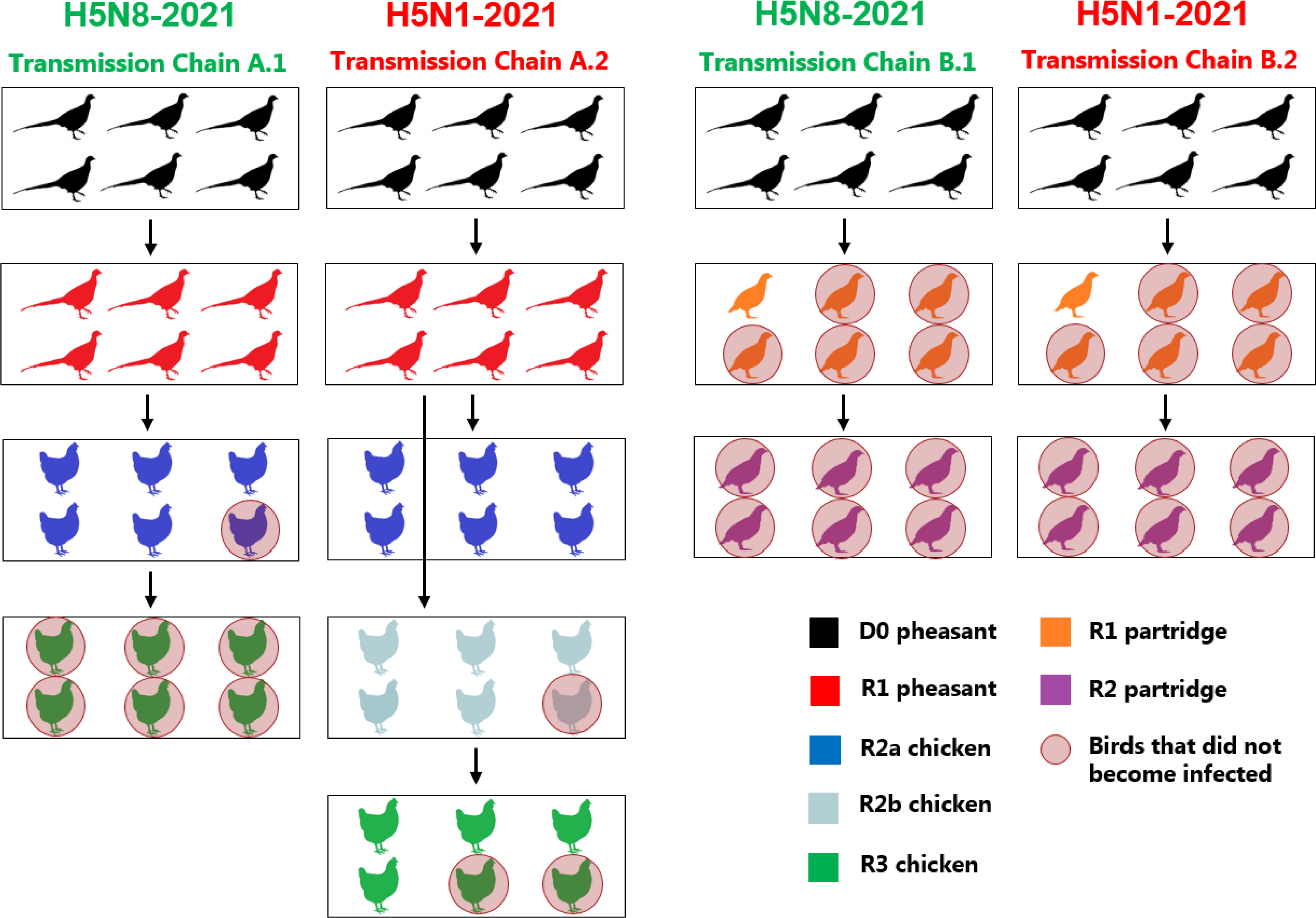
Schematic of the four intra- and inter-species transmission chains undertaken in this study. All transmission attempts commenced with the direct-inoculation of D0 (“donor”) pheasants with high doses of H5N8-2021 or H5N1-2021. The first groups of contact birds (R1 “recipients”) were introduced for co-housing at 1 dpi, with subsequent contacts (R-prefix groups) introduced as described in the Results, with only two successive groups of birds co-housed together at any time until study end. The blue chicken symbols represent R2 chickens in the H5N8-2021 transmission chain A1, and R2a chickens in transmission chain A2 (see Results for details). The circled birds represent birds that did not become infected following contact exposure, hence the originally-intended onward transmission of infection from partridges to chickens was not attempted because none of the R2 partridges became infected with either virus (transmission chains B.1 and B.2).

### Collection of clinical and environmental samples

Daily swabs (buccal and cloacal) were stored in 1 ml virus transport medium [38]. Tissue samples were collected *post-mortem*, as outlined in the Results. Environmental samples (faeces, straw/bedding and drinking water) were collected daily from each pen housing high-dose inoculated partridges in the MID_50_ experiment and in all four transmission chain experiments. Prior to total RNA extraction, processing was as previously for feathers [39] and for the other organs and environmental samples [40].

### Viral RNA detection: Matrix (M)-gene RRT-PCR

Total RNA was extracted robotically from supernatants of swab fluids, tissues and environmental samples by using a Qiagen (UK) BioRobot [41]. For each extracted specimen, 2ul of RNA was tested by M-gene RRT-PCR [42]. For quantitative interpretation, EID_50_ titrated H5N8-2021 and H5N1-2021 were extracted to provide vRNA which was diluted ten-fold to construct a standard curve, with Ct values converted to relative equivalent units (REUs) by correlation with the original EID_50_/ml values on the curves [43]. Specimens with Ct<36 were considered positive and Ct>36 were interpreted as negative [44]. An infected bird was identified by viral shedding if positive vRNA was detected in at least one swab at any given time. Swabs were collected from live birds, but occasional mortalities were further confirmed as being virus-associated by including *post-mortem* swab testing, as indicated in the Results and the relevant figure legends.

### Histology and immunohistochemistry

Tissue samples obtained at necropsy were fixed in 10% neutral-buffered formalin, processed and embedded into paraffin blocks for sectioning. Tissue sections were prepared and stained for haematoxylin and eosin (H&E), with virus-specific immunohistochemical (IHC) staining to detect influenza A virus nucleoprotein (NP) antigen by using an anti-NP monoclonal antibody [43]. Semi-quantitative scoring from 0 (absence of staining/lesion) to 4 (abundant staining/confluent lesions) was used for histopathological interpretation and IHC staining [45].

### Serology

Sera obtained from coagulated blood were heat-treated at 56° C for 30 minutes in a water bath, with pheasant and partridge sera pre-adsorbed as for duck sera [40]. Serum samples were tested for H5-specific antibodies by haemagglutination inhibition (HI), using four haemagglutination units of the homologous H5N8-2021 and H5N1-2021 antigens [36]. A reciprocal HI titre of 16 or above was considered as positive.

### Whole-genome sequencing

Whole genome sequences (WGSs) were obtained from viral inocula and deposited under accession numbers EPI2691150-EP2691157 (H5N8-2021) and EPI2691158-EPI291165 (H5N1-2021) [40]. WGS was also undertaken using RNA extracted from swabs and tissues, which required initial amplification with the primers MBTuni12R and MBTuni13 [46], as described [14]. Consensus sequences were generated using IRMA version 1.0.3 [47].

### Statistics

To quantify viral shedding from a given group of birds, the mean vRNA levels detected during shedding from the buccal and cloacal tracts were assessed by area under the curve (AUC) analysis. The AUCs were compared by the Mann-Whitney U test using GraphPad Prism 9.3.0 (GraphPad Software, Inc., San Diego, CA), with p<0.05 considered as statistically significant.

## Results

### H5N8-2021 infectivity (MID_50_), pathogenesis and mortality in pheasants

During this MID_50_ experiment, daily swabbing revealed that none of the pheasants directly inoculated with low-dose H5N8-2021 became infected. These pheasants were culled and bled at 8 dpi, with absence of H5 HI seroconversion confirming their uninfected status. All pheasants directly inoculated with the medium and high-doses shed H5N8-2021 (Figure 2a), and initially developed clinical signs which were largely mild or non-specific (dropped wings, huddling, ruffled feathers and lethargy). Prior to death, more severe neurological outcomes were frequently observed, namely loss of balance, tremors, torticollis and paralysis. In the medium-dose group, mean positive shedding was detected from 3 dpi and 83% (5/6) pheasants died by 10.25 dpi (Figure 2b). The remaining medium-dose pheasant was apparently healthy but infected at this time when it was culled to respect animal welfare (Figure 2b legend). A dose effect was apparent because the high-dose group experienced onset of mean positive shedding 1-day before the medium-dose group, with the high-dose group’s mortality being more rapid, with all six high-dose group pheasants dying by 4.25 dpi (Figure 2b). The MDT was 3.4 and 7.3 days in the high and medium-dose groups, respectively, with the pheasant H5N8-2021 MID_50_ being 10^3^ EID_50_ (Table 1).

**Fig. 2.**
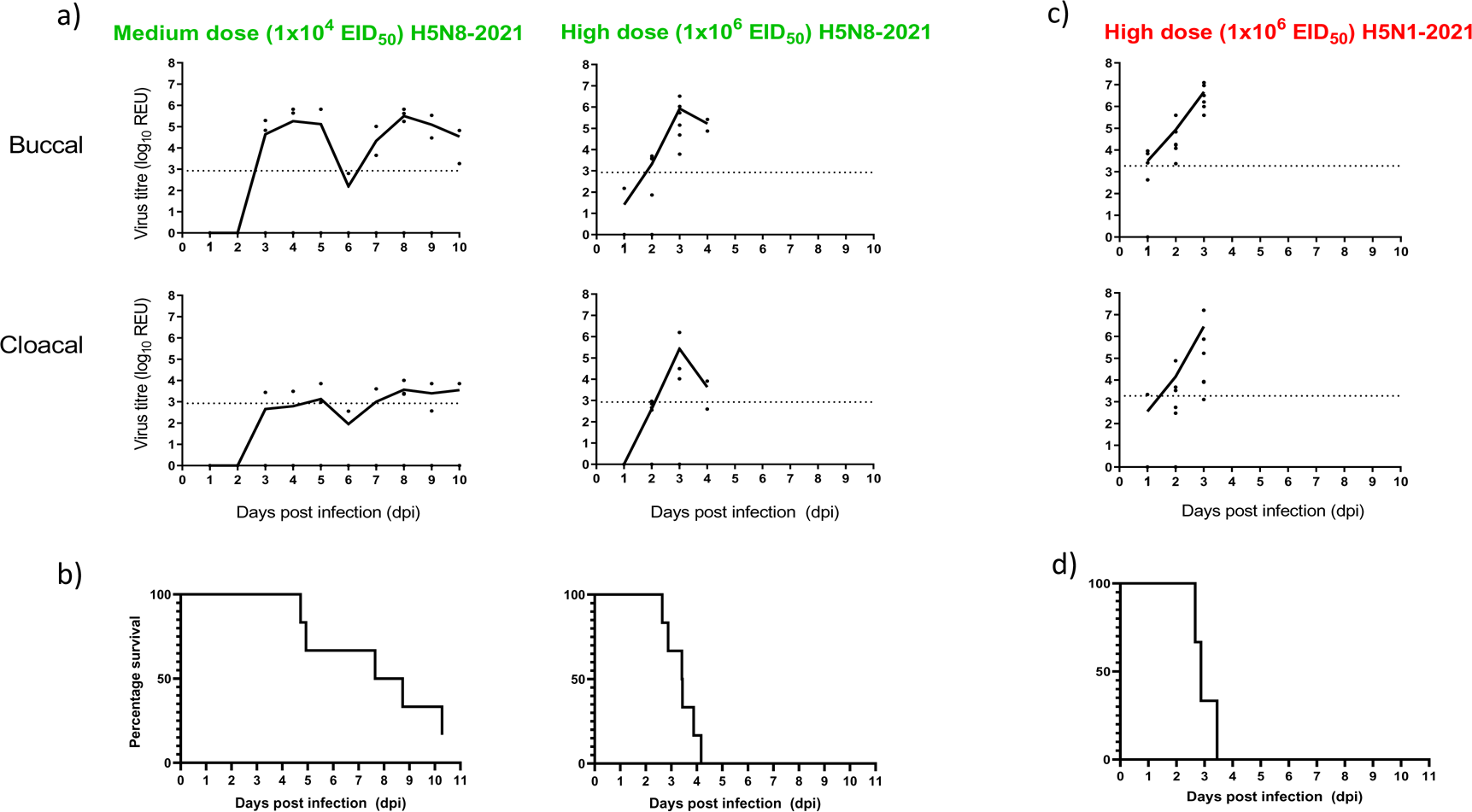
Viral shedding and survival of pheasants directly-infected with medium-dose and high-dose H5N8-2021, and high dose H5N1-2021. Shedding (buccal and cloacal, as indicated) is shown in (a) and (c) for H5N8-2021 and H5N1-2021 respectively. Symbols indicate shedding for individual pheasants, while continuous lines show the mean shedding (buccal and cloacal) for a given pheasant group. The positive cut-off for the M-gene RRT-PCR is indicated by the broken horizontal line, with the viral RNA levels denoted as log_10_ relative equivalent units (REUs). For the H5N8-2021 infected pheasants, AUC comparisons showed significantly greater buccal compared to cloacal shedding in the medium (p=0.043) and high-dose groups (p=0.041). For the H5N1-2021 infected pheasants (high-dose), there was a tendency for the buccal shedding to be greater than the cloacal shedding, but this was not considered as significant (p=0.093). (b) and (d) are Kaplan-Meier survival plots for pheasants directly infected with H5N8-2021 (medium and high doses) and H5N1-2021 (high dose), respectively. (b) includes one infected (but healthy) medium-dose pheasant which was culled at 10 dpi for welfare reasons which strongly advise against the prolonged housing of a single surviving bird, so the R1 survival over time is shown for the five virus (H5N1-2021)-associated deaths in this R1 group.

**Table 1.**
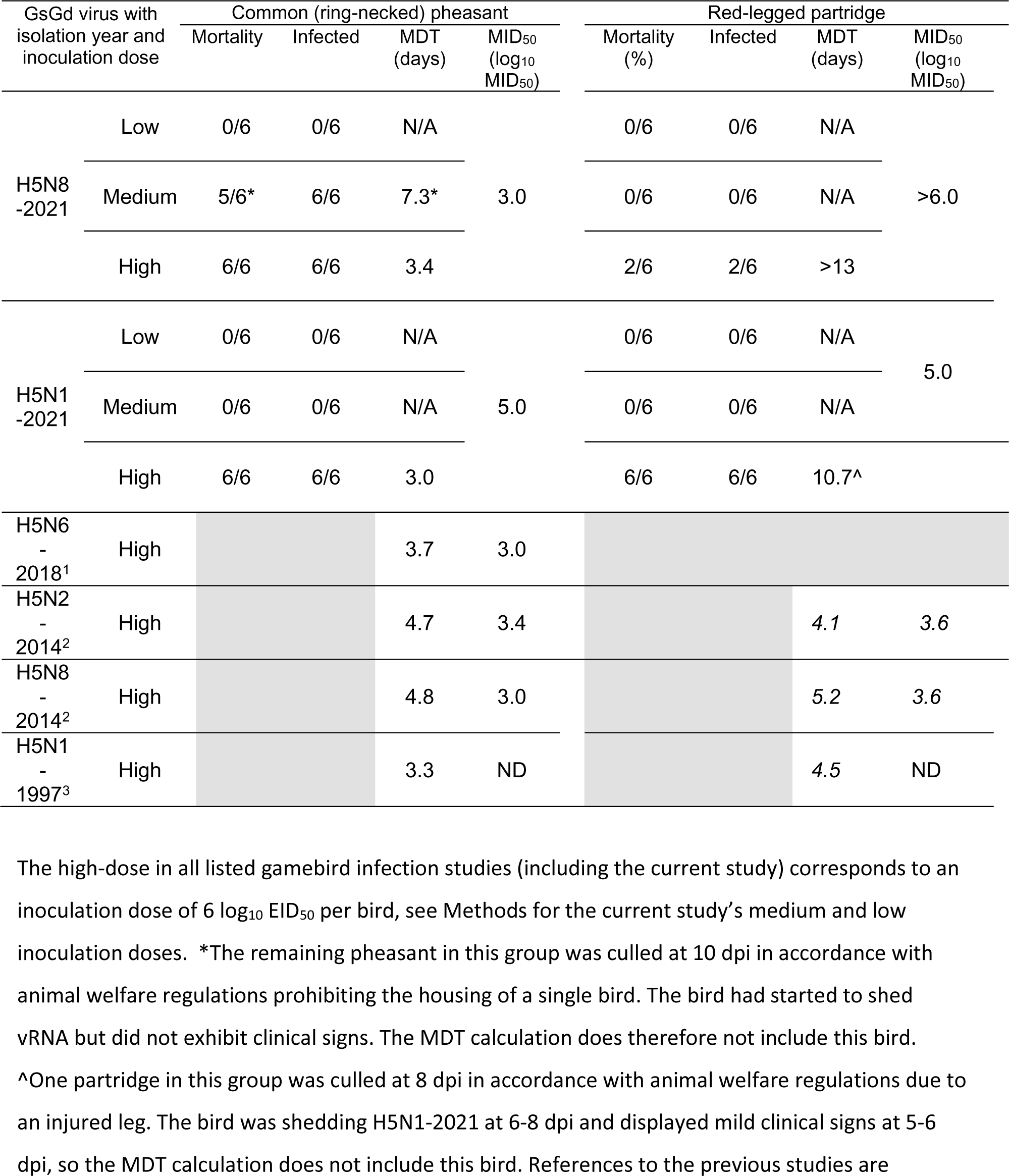
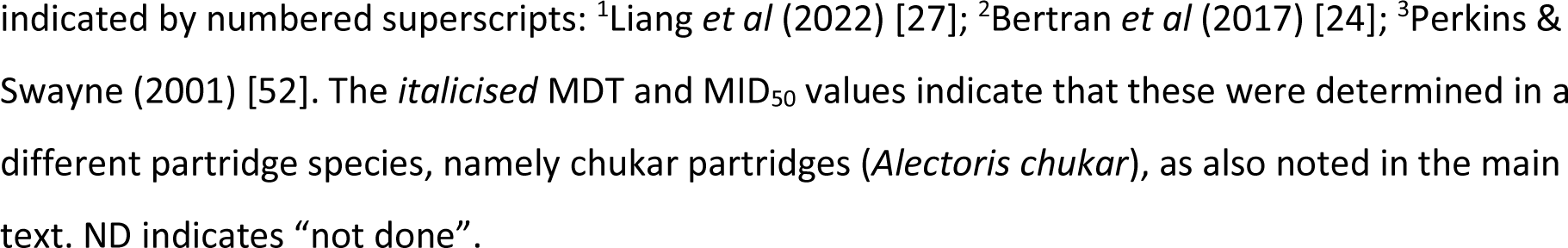
Infection and mortality details for pheasants and partridges following direct inoculation with the H5N8-2021 and H5N1-2021. Similar details are included from other H5Nx GsGd lineage gamebird infection studies.

| GsGd virus with isolation year and inoculation dose |  | Common (ring-necked) pheasant |  |  |  | Red-legged partridge |  |  |  |
| --- | --- | --- | --- | --- | --- | --- | --- | --- | --- |
|  |  | Mortality | Infected | MDT (days) | MID<sub>50</sub> (log<sub>10</sub> MID<sub>50</sub>) | Mortality (%) | Infected | MDT (days) | MID<sub>50</sub> (log<sub>10</sub> MID<sub>50</sub>) |
| H5N8 -2021 | Low | 0/6 | 0/6 | N/A |  | 0/6 | 0/6 | N/A |  |
|  | Medium | 5/6* | 6/6 | 7.3* | 3.0 | 0/6 | 0/6 | N/A | >6.0 |
|  | High | 6/6 | 6/6 | 3.4 |  | 2/6 | 2/6 | >13 |  |
| H5N1 -2021 | Low | 0/6 | 0/6 | N/A |  | 0/6 | 0/6 | N/A |  |
|  | Medium | 0/6 | 0/6 | N/A | 5.0 | 0/6 | 0/6 | N/A | 5.0 |
|  | High | 6/6 | 6/6 | 3.0 |  | 6/6 | 6/6 | 10.7^ |  |
| H5N6 - 2018<sup>1</sup> | High |  |  | 3.7 | 3.0 |  |  |  |  |
| H5N2 - 2014<sup>2</sup> | High |  |  | 4.7 | 3.4 |  |  | 4.1 | 3.6 |
| H5N8 - 2014<sup>2</sup> | High |  |  | 4.8 | 3.0 |  |  | 5.2 | 3.6 |
| H5N1 - 1997<sup>3</sup> | High |  |  | 3.3 | ND |  |  | 4.5 | ND |

The high-dose in all listed gamebird infection studies (including the current study) corresponds to an inoculation dose of 6 log<sub>10</sub> EID<sub>50</sub> per bird, see Methods for the current study's medium and low inoculation doses. \*The remaining pheasant in this group was culled at 10 dpi in accordance with animal welfare regulations prohibiting the housing of a single bird. The bird had started to shed vRNA but did not exhibit clinical signs. The MDT calculation does therefore not include this bird. ^One partridge in this group was culled at 8 dpi in accordance with animal welfare regulations due to an injured leg. The bird was shedding H5N1-2021 at 6-8 dpi and displayed mild clinical signs at 5-6 dpi, so the MDT calculation does not include this bird. References to the previous studies are
indicated by numbered superscripts: <sup>1</sup>Liang *et al* (2022) [27]; <sup>2</sup>Bertran *et al* (2017) [24]; <sup>3</sup>Perkins & Swayne (2001) [52]. The *italicised* MDT and MID<sub>50</sub> values indicate that these were determined in a different partridge species, namely chukar partridges (*Alectoris chukar*), as also noted in the main text. ND indicates “not done”.

The time-course pathogenesis experiment for pheasant H5N8-2021 infection revealed increasing viral load and numbers of infected tissues from 1-3 dpi (Figure S1a). Peak vRNA levels were detected by 3 dpi in all tissues, with the highest in the air sacs, heart, feathers and brain. Gross pathology at 2 dpi revealed enlarged and mottled spleens with mildly swollen kidneys, including splenic and pancreatic necrosis plus ascites by 3 dpi (Figure S2a). Five of the six pheasants were apparently healthy at the time of cull, with only pheasant #2024 displaying severe signs at euthanasia at 3 dpi. Histopathological changes were seen in the spleen and bursa from 2 dpi, spreading to other organs by 3 dpi, including the brain and pancreas (Figure S3a). Virus-specific IHC staining was observed in several tissues at 2 dpi, with the strongest staining in the feathers, heart, brain and pancreas at 3 dpi; and was overall strongest in the clinically affected pheasant #2024 (Figures S3b and S3e).

### H5N1-2021 infectivity (MID_50_), pathogenesis and mortality in pheasants

During this MID_50_ experiment, daily swabbing revealed that none of the pheasants directly inoculated with low or medium-dose H5N1-2021 became infected. These low and medium-dose pheasant groups were culled and bled at 8 and 11 dpi, respectively, with absence of H5 HI seroconversion confirming their uninfected status. The high-dose pheasant group shed mean positive levels of H5N1-2021 between 1-3 dpi (Figure 2c), by which time 100% (6/6) of these pheasants had died (Figure 2d), after experiencing severe disease which was the same as noted above for the H5N8-2021 infected pheasants. The pheasant H5N1-2021 MID_50_ was 10^5^ EID_50_, with a MDT of 3.0 days (Table 1).

The time-course pathogenesis experiment for H5N1-2021 pheasant infection revealed increasing viral load and numbers of infected tissues from 1-3 dpi (Figure S1b). Peak vRNA levels were again detected at 3 dpi in the air sacs, heart, feathers and brain, with generally higher H5N1-2021 vRNA levels attained in many tissues compared to H5N8-2021 infected pheasants (compare Figure S1a and b). The two pheasants which were found dead at 3 dpi displayed more pronounced pancreatic necrosis than the H5N8-2021 infected pheasants at the same time-point, plus splenic necrosis and ascites (Figure S2a). Histopathological changes and virus-specific staining due to H5N1-2021 infection progressed to many tissues between 2-3 dpi, with the strongest IHC staining observed in the heart, brain and pancreas (Figure S3a, b and e).

### H5N8-2021 infectivity (MID_50_), pathogenesis and mortality in partridges

During this MID_50_ experiment, daily swabbing testing revealed that none of the partridges directly-inoculated with low or medium-dose H5N8-2021 were shedding (data not shown) and remained apparently healthy throughout. These partridges were culled and bled at 10 dpi and 14 dpi, respectively, with absence of H5 HI seroconversion confirming their H5N8-2021 uninfected status. Similarly, none of the partridges inoculated with the H5N8-2021 high-dose shed virus (Figure 3a), but 33% (2/6) subsequently developed initial clinical signs at 5-6 dpi (dropped wings, huddling and ruffled feathers, diarrhoea) which became severe (loss of balance, tremors, torticollis, paralysis) and necessitated euthanasia at 6 and 8 dpi (Figure 3b). Viral RNA was detected across a broad range of tissues from both these partridges and included peak levels in the brain, while generally higher levels were detected in the enteric compared to the respiratory tract tissues (partridges #2142 (6 dpi) and #2140 (8 dpi), Figure S4a). Gross pathology changes were not observed in these two partridges, but mild histopathological changes occurred in the brain, heart, duodenum, colon and caecum (Figure S4c), with viral antigen also detected in these tissues (Figure S4d).

**Fig. 3.**
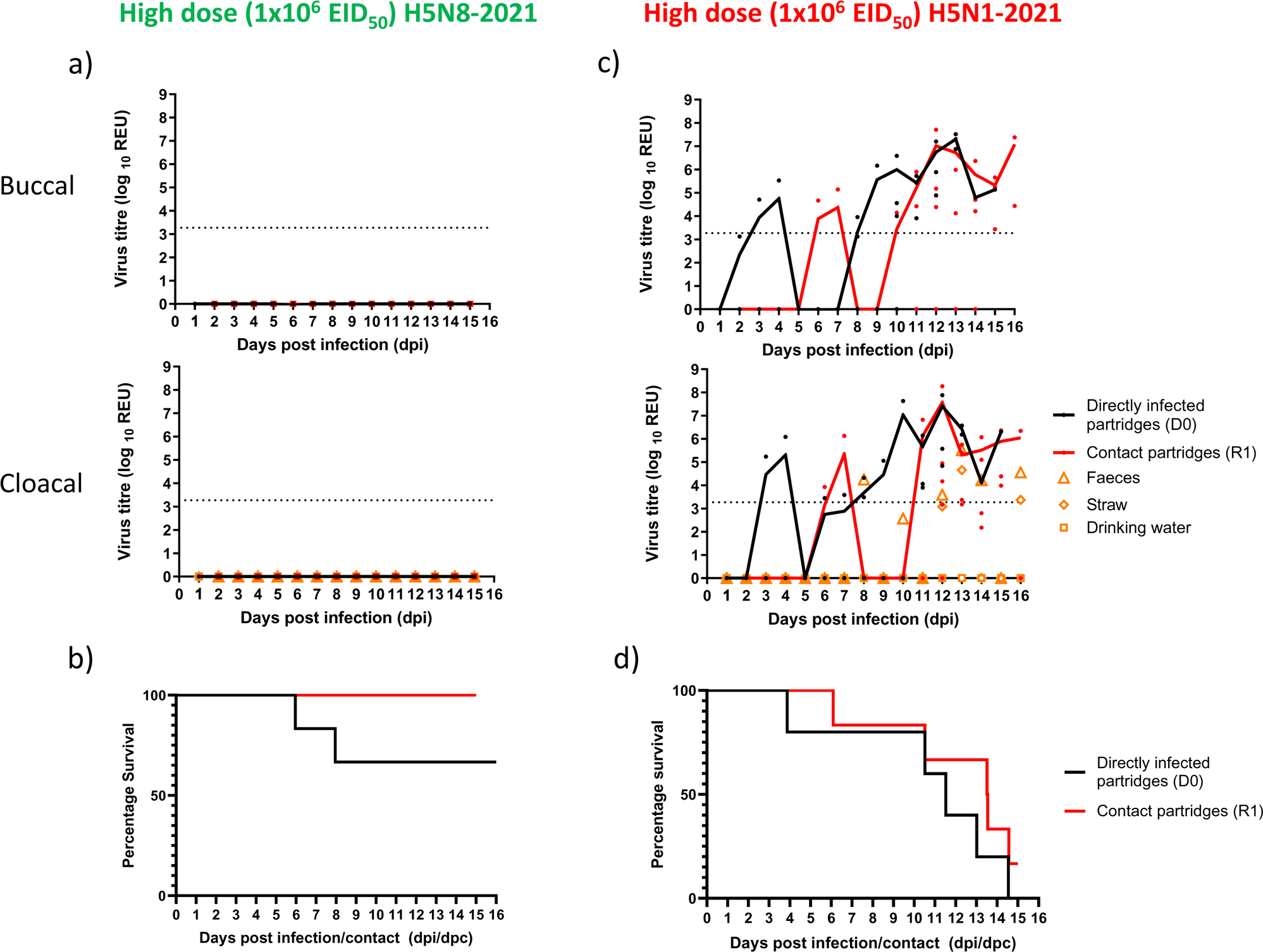
Viral shedding and survival of partridges infected with high doses of H5N8-2021 and H5N1-2021. Shedding (buccal and cloacal, as indicated) is shown in (a) and (c) for H5N8-2021 and H5N1-2021 respectively. Symbols indicate shedding for individual partridges, while continuous lines show the mean shedding (buccal and cloacal) for a given partridge group, with directly-infected (D0) and contact (R1) partridges indicated in black and red respectively. The cloacal shedding panels in (a) and (c) include vRNA levels in environmental samples, shown as orange symbols for faeces (triangles), straw/bedding (diamonds) and drinking water (squares), for both H5N8-2021 and H5N1-2021 partridge groups. The positive cut-off for the M-gene RRT-PCR is indicated by the broken horizontal line, with the viral RNA levels denoted as log_10_ relative equivalent units (REUs). AUC comparisons showed no statistically significant differences between the buccal and cloacal shedding for the H5N1-2021 infected D0 (p=0.937) and R1 (p=0.699) partridges. (b) and (d) are Kaplan-Meier survival plots for the partridge groups (both D0 and R1) following infection with H5N8-2021 and H5N1-2021, respectively. In (d), one directly-infected (D0) partridge required euthanasia due to leg injury, so the D0 survival plot is shown for five D0 partridges and the MDT also calculated for the same five D0 partridges (Table 1). (d) also includes one infected (but healthy) contact (R1) partridge which was also culled at 15 dpc for welfare reasons which strongly advise against prolonged housing of a single surviving bird, so the R1 survival over time is shown for the five virus (H5N1-2021)-associated deaths in this R1 group. Horizontal axes indicate days post-infection (dpi) for the directly-inoculated partridges and days post-contact (dpc) for the cohoused (from 1 dpi onwards) contact partridges.

At 1 dpi during this MID_50_ experiment, six contact (R1) partridges were co-housed with the high-dose H5N8-2021 inoculated (D0) partridges, but remained healthy and did not shed virus throughout, indicating no H5N8-2021 transmission (Figure 3a and b). All six R1 and the four surviving D0 partridges were confirmed as uninfected by H5 HI seronegative results at 15 dpi (14 dpc). Testing of daily environmental specimens from this partridge pen revealed total absence of environmental contamination by H5N8-2021 (Figure 3a). The MID_50_ in partridges infected with H5N8-2021 could not be specifically quantified, but was greater than 10^6^ EID_50_, with the MDT in the high-dose group being greater than 13 days (Table 1).

The time-course pathogenesis investigation focused on the early infection stage between 1-3 dpi and demonstrated absence of H5N8-2021 vRNA and viral antigen in all partridge tissues (Figures S1c, S3d and S3e). This observation affirmed the delayed H5N8-2021 pathogenesis in partridges compared to pheasants.

### H5N1-2021 infectivity (MID_50_), pathogenesis, and mortality in partridges

During this MID_50_ experiment, none of the partridges directly inoculated with low or medium-dose H5N1-2021 shed virus (data not shown), but remained healthy throughout, with absence of infection confirmed by negative serology. However, in contrast to the H5N8-2021 high-dose inoculated partridges (Figure 3a), all six D0 partridges inoculated with the H5N1-2021 high-dose demonstrated viral shedding from 3 dpi (Figure 3c). Clinical disease included conjunctivitis, dropped wings, subcutaneous oedema, lethargy, loss of balance and tremors, resulting in one requiring euthanasia and four being found dead during 4-15 dpi (Figure 3d). The sixth infected D0 partridge displayed mild signs (huddling and ruffled feathers) at 5-6 dpi, but required euthanasia due to an injured leg at 8 dpi. The partridge H5N1-2021 MID_50_ was 10^5^ EID_50_ with a MDT of 10.7 days (Table 1).

Six contact (R1) partridges were introduced at 1 dpi and all six (100%) became infected with H5N1-2021, with shedding first detected at 6 dpi (5 dpc; Figure 3c). Successful H5N1-2021 transmission in partridges clearly contrasted with H5N8-2021 partridge-to-partridge transmission failure, although five of six R1 partridges (83%) died by 14.6 dpc (Fig 3d), with the single remaining infected (but apparently healthy) partridge culled at 15 dpc. Unlike the environmental testing for the H5N8-2021 inoculated partridges (Figure 3a), the shedding partridges deposited virus-containing faeces and contaminated straw/bedding in the high-dose H5N1-2021 pen, with vRNA detectable from 8-16 dpi (Figure 3c).

Testing of tissues from H5N1-2021 partridge deaths from 4 dpi onwards revealed vRNA in the respiratory and enteric tracts, plus the brain, heart, spleen, pancreas and kidney (Figure S4b). Stronger H5N1-2021 histopathological changes were observed in the pancreas than for H5N8-2021 partridge infections (Figure S4c), with stronger viral antigen in the feathers, pancreas and spleen (Figures S2c and S4d). During the time-course pathogenesis investigation, H5N1-2021 again differed from H5N8-2021 with vRNA already detected at 3 dpi in the heart, liver, pancreas, kidney and enteric tissues from one partridge (#2163) (compare Figure S1c and d). No histopathological changes occurred in this partridge, but viral antigen was detected in the feathers, with weaker staining in the proventriculus and kidney (Figures S3c, d and S3e). While pathogenesis was delayed in H5N1-2021 infected partridges compared to pheasants, pathogenesis onset and systemic viral spread in partridges was more rapid for H5N1-2021 than for H5N8-2021.

### Transmission chain A.1: H5N8-2021 pheasant transmission and onward transmission to chickens

To initiate this transmission chain, six D0 pheasants were H5N8-2021 infected, with transmission to R1 pheasants affirmed by shedding from 3 dpi (2 dpc; Figure 4a). By 4 dpi, all six infected D0 pheasants had died (Figure 4b), with six R2 chickens introduced at 4 dpi for co-housing with the R1 pheasants. By 4 dpi, all R1 pheasants were infected with 83% (5/6) mortality by 6 dpi. Inclusion of R1 contact pheasants represented a natural route of H5N8-2021 acquisition, where AUC comparisons showed higher buccal compared to cloacal shedding in the R1 pheasants (p=0.026). Viral RNA in faeces and drinking water was detected between 4-7 dpi (Figure 4a). Shedding from R2 chickens was detected at 2 dpc (6 dpi from initial D0 pheasant inoculation), with rapid mortality following where 83% (5/6) died by 4 dpc. Due to in the first R2 chicken mortality at 6 dpi (2 dpc), the remaining R1 pheasant (infected, but healthy) was culled and removed to allow the transmission chain to continue. R3 chickens were introduced at 6 dpi for co-housing with R2 chickens to attempt onward chicken-to-chicken transmission. After R3 chicken introduction (6 dpi), only 66.6% (2/3) of the remaining R2 chickens were shedding H5N8-2021 until death at 7 and 8 dpi (3 and 4 dpc, Figure 4b). Gross pathology of the chicken mortalities showed mild hydropericardium and epicardial petechiae, with necrosis and strong virus-specific IHC labeling in the lung and spleen, plus IHC staining in the heart (Figure S2a and b). There was no H5N8-2021 shedding in the last surviving R2 chicken, or in any of the six R3 chickens throughout. These seven surviving chickens did not seroconvert by 15 dpi, which confirmed their uninfected status.

**Fig. 4.**
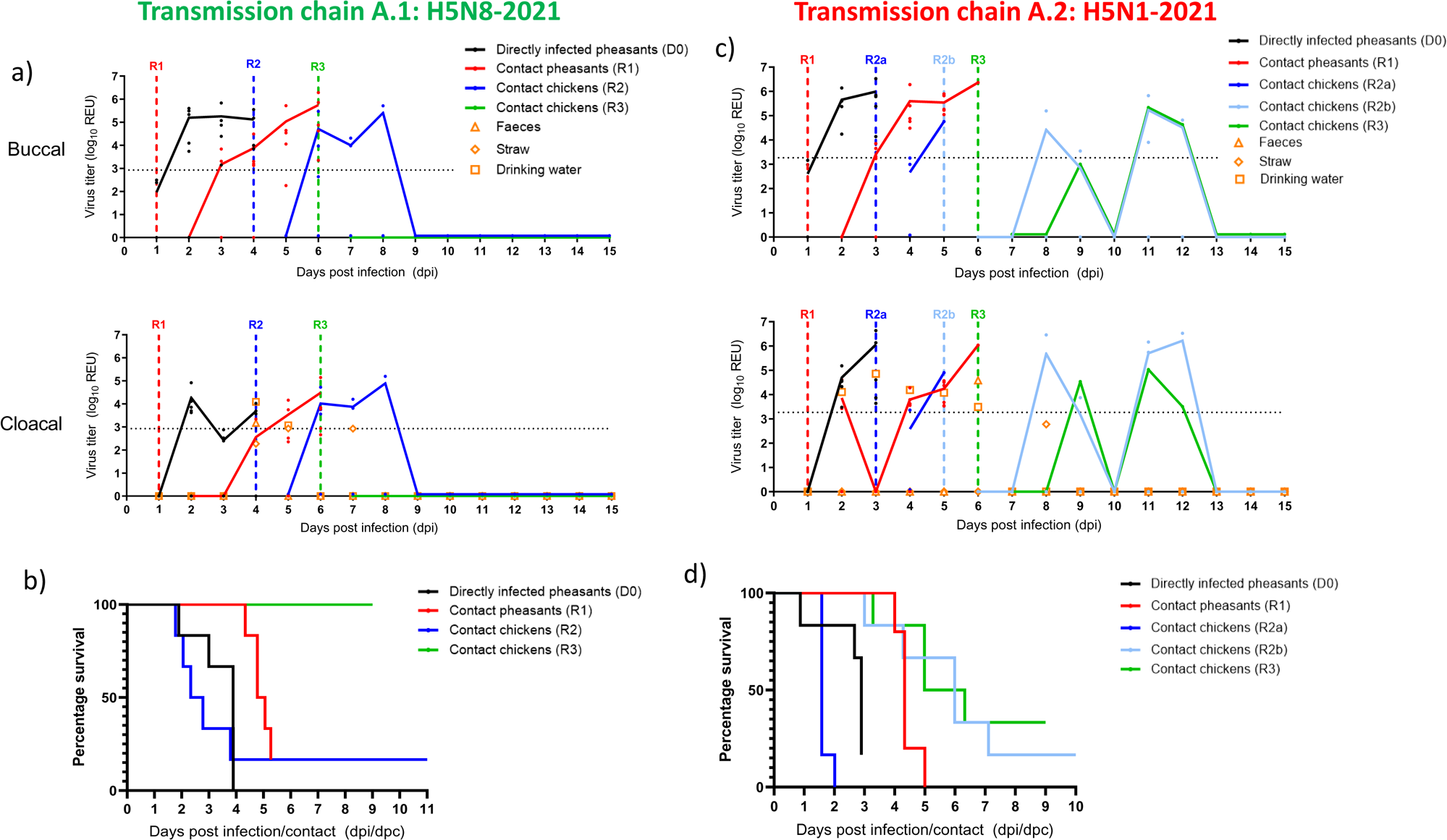
Viral shedding and survival of H5N8-2021 (a-b) and H5N1-2021 (c-d) intra- and inter-transmission among pheasants and chickens. (a) and (c) Individual (symbols) and mean (lines) indicate buccal and cloacal shedding. Viral RNA levels are indicated for environmental samples as in Figure 3. The positive cut-off for the M-gene RRT-PCR is indicated by the broken horizontal line, with the viral RNA levels denoted as log_10_ relative equivalent units (REUs). Vertical broken lines indicate the introduction time-points of the contact groups. (b) and (d) Kaplan-Meier survival plots of survival of pheasants and chickens following H5N8-2021 and H5N1-2021 intra- and inter-species transmission. Horizontal axes indicate days post-infection (dpi) for the D0 pheasants and days post-contact (dpc) for the contact (R-prefix) bird groups introduced for cohousing. In (d), one D0 pheasant was removed to enable continuation of the transmission chain hence only five mortalities are indicated (see Results text), and one R1 pheasant was culled due to a leg injury and is not included in the graph which presents only virus-related deaths. The R2b and R3 chicken mortalities shown in (d) were confirmed as being due to H5N1-2021 infection on the basis of virus positive swabs collected from the carcasses, as shown in (c).

### Transmission chain A.2: H5N1-2021 pheasant transmission and onward transmission to chickens

To initiate this transmission chain, six D0 pheasants were H5N1-2021 infected, with successful transmission to R1 pheasants affirmed by shedding from 2 dpi (1 dpc), with all six (100%) R1 pheasants infected by 4 dpi (3 dpc) (Figure 4c). Inclusion of R1 contact pheasants represented a natural route of H5N1-2021 acquisition, where AUC comparisons showed higher buccal compared to cloacal shedding in the R1 pheasants (p=0.026). At 3 dpi, all D0 pheasants were infected with 83% (5/6) mortality, so six chickens were introduced (R2a) (Figure 4c and d). At 4 dpi (1 dpc), the first shedding occurred in 33% (2/6) of the R2a chickens. At 4.5 dpi (1.5 dpc), rapid R2a chicken mortality followed with 83% (n=5/6) dying, and the final R2a chicken died at 5 dpi (2 dpc). Gross pathology of R2a chicken mortalities revealed a high degree of hydropericardium, epicardial petechiae and ascites, with necrosis and strong virus-specific IHC labeling in lung and spleen, plus IHC staining in the heart, (Figure S2a and c).

By 5 dpi (4 dpc), only one R1 pheasant had died, and another was culled for welfare reasons due to a leg injury. The remaining four live R1 pheasants were H5N1-2021 infected, so an additional group of six chickens were introduced (R2b) at 5 dpi to replace the six R2a mortalities. Subsequently, on the same day, three of the remaining four R1 pheasants were euthanized following the onset of severe clinical disease, including torticollis and paralysis. The last R1 pheasant died at 6 dpi (5 dpc). The introduction of R2b chickens was at a late stage of R1 pheasant infection, and they were only co-housed for a maximum of one day before the final R1 pheasant death, so R3 chickens were introduced at 6 dpi. Mortality affected 83% (5/6) of the R2b chickens between 8-12 dpi (3-7 dpc), but no shedding occurred on the days prior to death, with infection only confirmed at terminal swabbing, while one R2b chicken remained alive and uninfected (Figure 4c and d). Onward transmission of H5N1-2021 from R2b chickens to R3 chickens was observed with 67% (4/6) becoming infected and dying (Figure 4d). Again, the R3 chickens exhibited a similar shedding pattern which was limited to the day of death, whilst two R3 chickens remained uninfected. The uninfected status of the three surviving chickens was confirmed by absence of seroconversion at 15 dpi.

During H5N1-2021 transmission among chickens (6-12 dpi), the final instance of positive environmental contamination was detected at 6 dpi when the final R1 pheasant died, with only a single subsequent instance of sub-threshold (negative) vRNA in the straw/bedding at 8 dpi (Figure 4c). H5N1-2021 vRNA was otherwise detected earlier in the drinking water (2-6) dpi and in faeces at 6 dpi.

### Transmission chain B.1: H5N8-2021 pheasant-to-partridge transmission

To attempt this transmission chain between the two gamebird species, six D0 pheasants were H5N8-2021 infected with R1 partridges introduced at 1 dpi. The six infected D0 pheasants all died by 4 dpi (Figure 5a and b), so R2 partridges were introduced at this time-point. No H5N8-2021 shedding was detected from the R1 or R2 partridges throughout the experiment to its conclusion at 16 dpi. However, at 9 dpi (8 dpc) one R1 partridge was euthanized due to sudden onset of severe disease which included diarrhoea, loss of balance, tremors and torticollis. H5N8-2021 vRNA was detected in a broad range of tissues from this partridge (#2106; Figure S4a), with accompanying histopathological changes in the brain, bursa, duodenum and colon (Figure S4c). Viral antigen was detected in the lung, duodenum, colon, caecum and brain (Figure S4d). There was no seroconversion in the remaining five R1 partridges or all six surviving R2 partridges by 15 dpi, affirming their uninfected status. H5N8-2021 vRNA was detected in drinking water, faeces and straw/bedding (2-6 dpi; Figure 5a). Absence of H5N8-2021 shedding from any contact partridges suggested that this environmental contamination originated from the infected D0 pheasants.

**Fig. 5.**
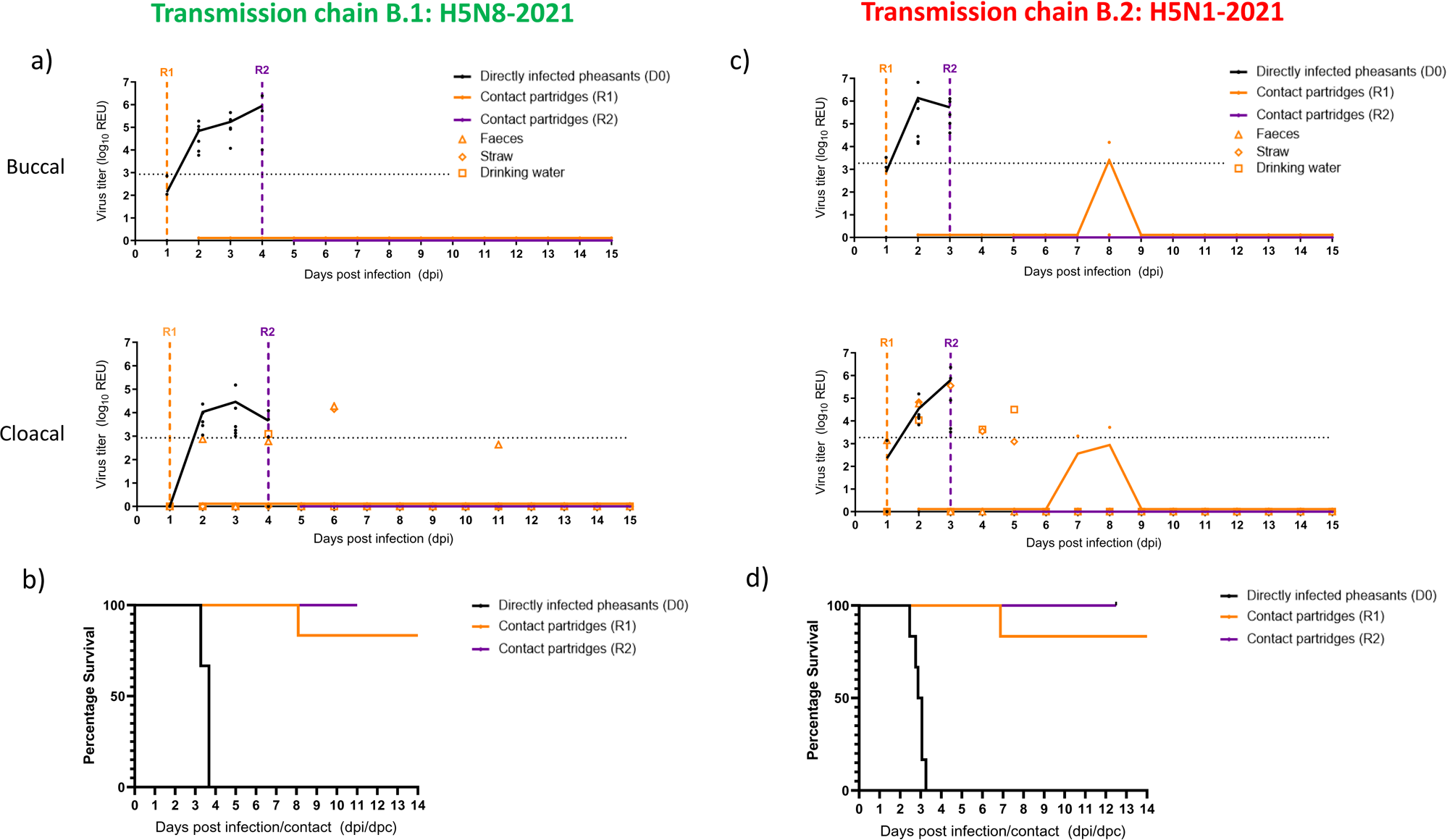
Viral shedding and survival of (a-b) H5N8-2021 and (c-d) H5N1-2021 during intra- and inter-species transmission attempts among pheasants and partridges. (a) and (c) Individual (symbols) and mean (line) buccal and cloacal shedding. The three H5N1-2021 positive swabs obtained from the R1 partridge (#2111) included buccal and cloacal swabs collected at *post-mortem* at 8 dpi (7 dpc). Viral RNA levels are indicated for environmental samples as in Figure 3. Vertical broken lines indicate the introduction time-points of the contact groups. The positive cut-off for the M-gene RRT-PCR is indicated by the broken horizontal line, with the viral RNA levels denoted as log_10_ relative equivalent units (REUs). (b) and (d) Kaplan-Meier survival plots of survival of pheasants and partridges following H5N8-2021 and H5N1-2021 intra- and inter-species transmission. Horizontal axis indicates days post-infection (dpi) for the D0 pheasants and days post-contact (dpc) for the contact (R-prefix) partridges introduced for cohousing.

### Transmission chain B.2: H5N1-2021 pheasant-to-partridge transmission

To initiate this transmission chain, six D0 pheasants were H5N1-2021 infected, with all dying rapidly by 3.25 dpi (Figure 5c and d). R1 partridges were introduced at 1 dpi, followed by R2 partridges at 3 dpi to replace the D0 pheasants. A single R1 partridge (#2111) shed H5N1-2021 at one time-point (7 dpi [6 dpc]) and died (8 dpi [7 dpc]). This inefficient inter-species acquisition of H5N1-2021 by R1 partridges from D0 pheasants contrasted with the efficient intra-species transmission of H5N1-2021 from D0 to R1 partridges (Figure 3c). The *post-mortem* swabs from partridge #2111 were also vRNA positive (Figure 5c), together with many tissues (Figure S4b). Histopathological changes with viral antigen were detected in the heart, pancreas, duodenum, and colon (Figure S4c and d) of partridge #2111. H5N1-2021 shedding was not detected in the remaining five R1 and six R2 partridges, with none seroconverting by 15 dpi, affirming their uninfected status. H5N1-2021 vRNA was detected in drinking water, faeces and straw/bedding (2-5 dpi; Figure 5c).

### Genetic polymorphisms

Thirty-three WGSs were obtained from clinical specimens from each species from the *in vivo* experiments (13 and 20 WGSs following H5N8-2021 and H5N1-2021 infection, respectively; Table S1). Consensus sequences were compared with the respective inoculum sequence to identify emerging genetic polymorphisms, with four amino acid changes identified following H5N8-2021 infection (Table S1a). Direct H5N8-2021 inoculation (MID_50_ experiment, medium-dose; Figure 2) of a pheasant resulted in the PB2-W49G polymorphism. The H5N8-2021 transmission chains (Figures 4a and 5a) included NP-M374I emergence after transmission to a R1 pheasant and R2 chicken, with PA-I348V and NP-P83H identified in a R1 partridge.

During H5N1-2021 infection and transmission, three amino acid polymorphisms were identified (Table S1b), including PB2-G459E (R3 chicken) which did not occur earlier at the pheasant (D0 and R1) and chicken (R2a and R2b) transmission stages (Figure 4c). Direct H5N1-2021 inoculation (MID_50_ experiment, high-dose; Figure 3c) resulted in two NP changes (N247S and V286F) in two D0 and three R1 partridges.

## Discussion

The continued circulation of H5Nx clade 2.3.4.4b HPAIVs [12,48] necessitates better understanding of its pathogenesis and transmission in different bird species. We investigated the susceptibility of ring-necked pheasants and red-legged partridges to clade 2.3.4.4b HPAIVs of the H5N8 and H5N1 subtypes, isolated from farmed UK pheasants during early 2021. Novel outcomes included important differences between the H5N8-2021 and H5N1-2021 infections, which differed between both gamebird species. In assessing gamebirds as a bridging host, pheasants were more likely than partridges to transmit onwards to chickens, with H5N1-2021 representing a greater risk than H5N8-2021 (see Figure 1 which summarises the transmission outcomes).

High pheasant mortality was observed among all the H5N8-2021 and H5N1-2021 infected pheasants and may well have registered 100% in all groups, but for the pre-emptive culls prompted by welfare concerns. This observation resembled the high mortalities caused by recent (isolated since autumn 2020) UK H5N8 and H5N1 clade 2.3.4.4 HPAIVs in chickens [45,49]. H5N8-2021 and H5N1-2021 pheasant infections caused severe neurological signs prior to death. Gross lesions included pancreatic necrosis, as observed during the pheasant outbreaks where both viruses originated, and also during earlier and subsequent UK H5Nx HPAIV pheasant cases [20,50,51]. Both H5N8-2021 and H5N1-2021 displayed similar systemic tropism, but stronger H5N1-2021 pheasant pathogenesis was evident in higher vRNA levels across most tissues compared to H5N8-2021, and was also reflected in pheasant MDT differences (Table 1). Previous H5Nx GsGd lineage investigations in pheasants included European (H5N6-2018) and North American (H5N8 and H5N2, December 2014 isolates) clade 2.3.4.4 HPAIVs, which also disseminated systemically and caused high mortality [24,27,52].

H5N8-2021 pheasant infectivity resembled the other investigated H5Nx clade 2.3.4.4 HPAIVs, except for H5N1-2021 which had the highest MID_50_ (Table 1). Despite the H5N8-2021 and H5N1-2021 infectivity differences following pheasant direct-inoculation, both viruses transmitted efficiently among pheasants. Cohousing provided a more natural route for spread within a flock, which remained uninhibited by the ongoing pheasant mortality. It is plausible that viral environmental contamination contributed to efficient pheasant-to-pheasant transmission of both viruses, with H5N8-2021 and H5N1-2021 vRNA detected in faeces and straw/bedding, which followed cloacal shedding. Drinking water may have been contaminated by pheasant buccal shedding, although faecal (cloacal) contamination cannot be excluded (Figures 4 and 5). However, R1 pheasant buccal shedding was significantly greater than cloacal for both viruses. The drinking water was refreshed daily, hence vRNA detection in this matrix represented contamination during the preceding 24 hours, while detection in the solid matrices may represent earlier contamination since the initial shedding onset. There was no significant difference when pheasant H5N8-2021 and H5N1-2021 shedding were compared separately for each tract (data not shown). Efficient spread of both H5N8-2021 and H5N1-2021 among pheasants resembled the sustained transmission of UK-origin H5Nx clade 2.3.4.4 HPAIVs among ducks [35], where viral environmental contamination followed shedding and coincided with efficient onward transmission [40,45].

H5N1-2021 acquisition by chickens at the R3 contact stage was successful, but not accompanied by viral environmental contamination, hence chicken-to-chicken transmission was more likely due to close contact, as suggested elsewhere [49]. It is uncertain whether other factors linked to prior transmission among pheasants may have assisted the relatively successful onward transmission of H5N1-2020 among chickens, but onward spread of H5N8-2020 among chickens was relatively less effective. Other clade 2.3.4.4 chicken-to-chicken transmission attempts featured similar numbers of prior infected chickens, but resulted in zero or inefficient transmission [35,45,49,53].

The reduced susceptibility of H5N8-2021 partridge infection compared to pheasants was another novel finding. Interestingly, H5N8-2021 partridge infection was characterised by a total absence of viral shedding, whether after direct high-dose inoculation (Figure 3a), or during cohousing with D0 infected pheasants during H5N8-2021 attempted transmission among gamebirds (Figure 5a). H5N8-2021 infection was suspected following deaths in three partridges (Figures 3b and 5b), which was confirmed by H5N8-2021 tropism in multiple organs, with peak vRNA levels in the brain (Figure S4a and d). This manifestation of H5N8-2021 partridge infection contrasted with pheasant H5N8-2021 shedding, MID_50_ differences also apparent between both species for H5N8-2021 (Table 1). The partridge H5N8-2021 MDT was also longer, and reflected the slower H5N8-2021 pathogenesis onset in partridges compared to pheasants (Figure S1a and c). Importantly, the absence of H5N8-2021 shedding also prevented partridge-to-partridge transmission.

The contrast between H5N8-2021 (non-shedding) and H5N1-2021 (shedding) partridge infections was an important and novel finding, with successful partridge-to-partridge transmission only occurring for H5N1-2021 (Figure 3c). High mortality affected the H5N1-2021 infected partridges which, again, may have attained 100% but for welfare considerations (Figure 3d). Successful intra-species acquisition of H5N1-2021 by R1 partridges showed that this subtype represented a greater transmission risk in partridges than H5N8-2021. However, when modelling inter-species pheasant-to-partridge transmission, only one of six (17%) R1 partridges became infected and died (Figure 5d). Weak H5N1-2021 shedding from this R1 partridge appeared to be insufficient to transmit infection to R2 partridges, which all remained uninfected (Figure 5c). Environmental H5N1-2021 contamination may have also contributed to the efficient partridge-to-partridge transmission, and also to the less efficient pheasant-to-partridge transmission. Systemic tropism of H5N1-2021 infections in partridges affected a greater number of organs and attained higher levels than H5N8-2021 infection in this host (Figure S4), which, as in the case of pheasants, again underlined a more aggressive partridge pathogenesis for H5N1-2021 which included a shorter MDT. Inoculation of chukar partridges and ring-necked (common) pheasants with the North American H5N8-2014 and H5N2-2014 HPAIVs revealed similar infectivity for both viruses in both hosts, including shedding, systemic dissemination and high mortality (Table 1). However, H5N2-2014 appeared to transmit more readily to contact chukar partridges than H5N8-2014 [24] which again reflected how different clade 2.3.4.4 genotypes differ following partridge infection. We showed that early-stage replication in red-legged partridges differed between H5N8-2021 and H5N1-2021, as evidenced by initial systemic spread in one partridge at 3 dpi following direct H5N1-2021 inoculation, but none occurred at the same stage following similar high-dose H5N8-2021 partridge inoculation (Figure S1c and d). By combining this H5N1-2021 partridge pathogenesis observation with the timing of shedding from high-dose inoculated partridges (Figure 3c), H5N1-2021 disseminated systemically within partridges before shedding. Compared to pheasants, both viral infections demonstrated longer incubation in partridges prior to the observed mortalities.

Onward spread among partridges occurred relatively inefficiently for H5N1-2021 compared to that in pheasants, at least when modelled on an experimental scale, while no detectable H5N8-2021 partridge shedding would explain the absence of any partridge infections at mixed gamebird premises in the UK during the 2020-2021 H5N8 epizootic. During the earlier H5N8 clade 2.3.4.4 European epizootic in early 2017, UK pheasant outbreaks included readily observable mortality, although a proportion of the surviving field-infected pheasants displayed successful seroconversion [20]. Despite some possible resolution of H5N8-2017 infection in these farmed pheasants, the clear observation of suspect clinical signs (including neurological changes and mortality) prompted disease investigation. In summary, pheasants may be considered an appropriate indicator host at mixed gamebird premises for passive surveillance, based on the more rapid onset of clinical signs and mortality. Our H5N8-2021 and H5N1-2021 findings showed that any incursion into a mixed gamebird premises is more likely to initially manifest and spread efficiently among pheasants, before causing any subsequent clinical outcomes in partridges.

The H5N8-2021 and H5N1-2021 isolates in this study originated from pheasant outbreaks, hence their efficient spread within this host may reflect an existing prior adaptation, and /or improved host adaptation acquired during the pheasant outbreaks. It is speculated that the more successful H5N1-2021 onward transmission among chickens in the current study may have been aided by this isolate being of pheasant (galliforme)-outbreak origin, possibly assisted by emergence of the PB2 G459E polymorphism, as detected in one R3 chicken. No chickens were available to investigate whether H5N1-2021 transmission could be sustained further to an R4 chicken group. It was previously suggested that the occasional instances of H5N6-2017 clade 2.3.4.4 acquisition by contact chickens from infected ducks included polymerase gene changes, which may have reflected the virus attempting to become more adapted to chickens [40]. During the transmission attempts from H5N8-2021 infected pheasants, efficient acquisition occurred only in the first chicken contacts (R2), with zero transmission to the subsequent introduced R3 chicken group. No polymerase gene polymorphisms emerged during this H5N8-2021 transmission chain, although an NP M374I change occurred in one R1 pheasant and one R2 chicken.

During the extensive UK H5Nx clade 2.3.4.4 incursions since autumn 2020, continuing wild bird surveillance has rarely discovered partridge mortalities [54]. The current study may explain this observation by the relatively low infectivity of H5N8-2021 and H5N1-2021, with wild partridges featuring a long subclinical incubation, as noted already. Partridges may be able to mount a generally successful host response to infection, whereby H5N8-2021 was restricted to systemic infection with no shedding and subsequent transmission. Interestingly, following partridge H5N1-2021 inoculation, two genetic polymorphisms emerged in the NP gene, N247S and V286F, and both were also identified in the cohoused contact partridges. The more likely potential for H5N1-2021 than H5N8-2021 to spread among partridges has been noted, and while the significance of these NP mutations is unclear, they may indicate further adaptation to partridges. However, during 2022, no H5Nx clade 2.3.4.4 HPAIV outbreaks were reported at UK partridge farms, despite the continuing infection pressure due to the extensive H5N1 epizootic [55].

To conclude, our findings are of important interest from the perspective of understanding avian species susceptibility to the H5Nx clade 2.3.4.4 HPAIVs, particularly in view of the H5N1 subtype’s subsequent dominance in Europe and spread to the Americas [12,48,56,57]. The gamebird sector will continue to pose challenges, where it is important to assess the risks of infection spread which will be informed by the comparative resilience to infection in the relevant species. This knowledge will guide gamebird surveillance strategies. In comparing the infection dynamics of both H5N8 and H5N1 viruses in gamebirds, we demonstrated H5N1-2021 transmission from pheasants to chickens, and also onwards among chickens.

Furthermore, H5N1-2021 intraspecies spread between partridges was also observed, which did not occur for H5N8-2021. These fitness differences could therefore explain the continued circulation of H5Nx clade 2.3.4.4 HPAIVs, especially H5N1, in Europe. The persisting threat of clade 2.3.4.4 epizootics and continuing evolution of these viruses necessitates better understanding of the current and future circulating subtypes and genotypes of these HPAIVs, their diverse host interactions and associated infection dynamics.

## Acknowledgements

The authors would like to acknowledge the Animal Sciences staff at APHA for the collection of samples and assisting with the *in vivo* experiments. We acknowledge the authors, originating and submitting laboratories of the sequences from GISAID EpiFlu^TM^ Database on which this research is based. All submitters of the data may be contacted directly via the GISAID website (www.gisaid.org).

## Author contributions

All authors have read and agree to the published version of the manuscript. Conceptualization and funding, CKH, HS, IHB, LEL, ACB, MJS. Sample collection, processing, testing and experimentation, AHS, YL, CJW, SST, FZXL, AN, PS, DS. Computational data analysis, AHS, YL. Writing-original draft and preparation, AHS, YL, MF. Writing – review and editing, all authors.

