## Supplementary material for "Transmission dynamics and pathogenesis differ between pheasants and partridges infected with clade 2.3.4.4b H5N8 and H5N1 high-pathogenicity avian influenza viruses": Suppl. Figures S1-S4

### Slide 1
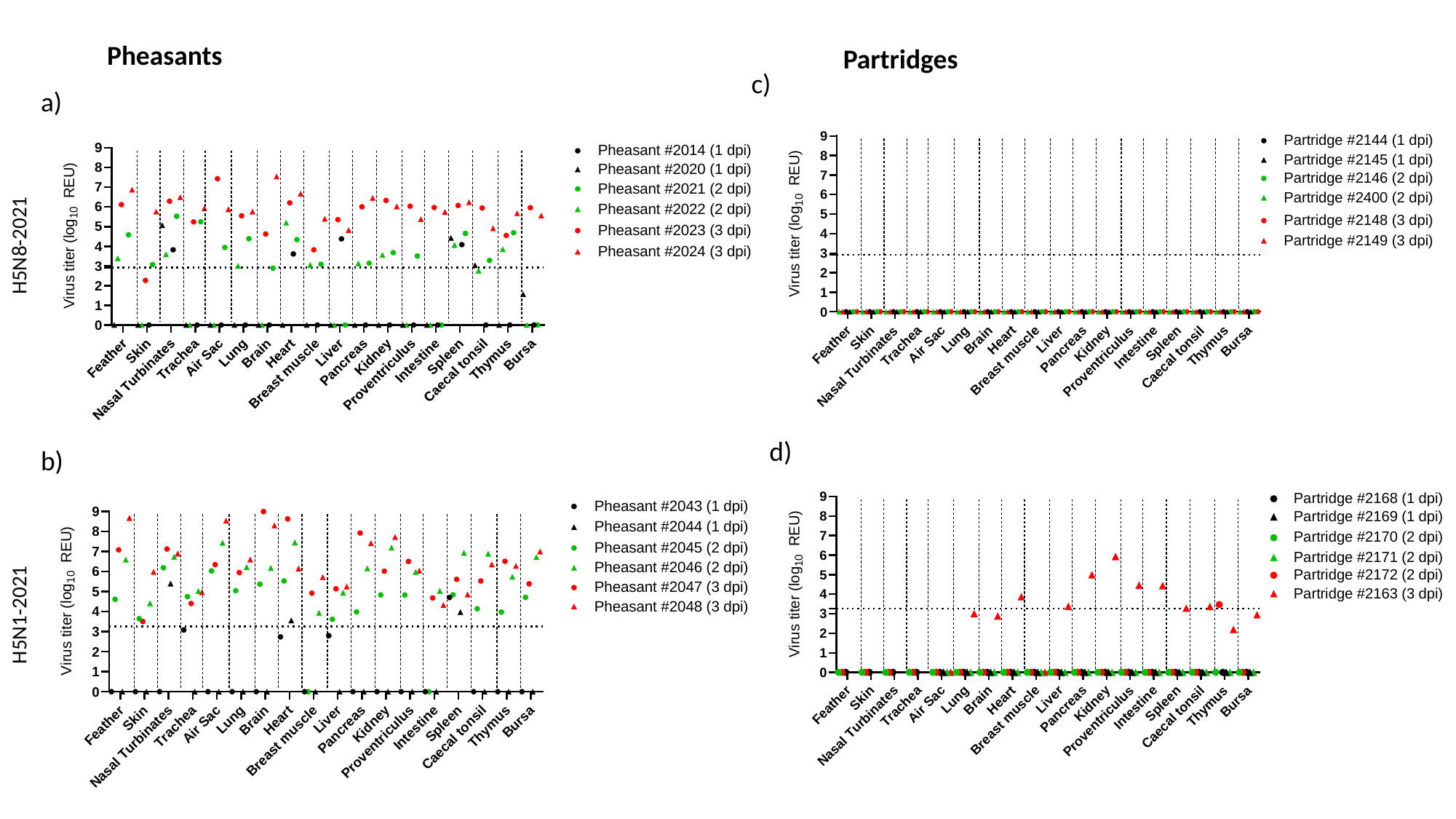

Pheasants
Partridges
c)
a)
H5N8-2021
d)
b)
H5N1-2021

### Slide 2
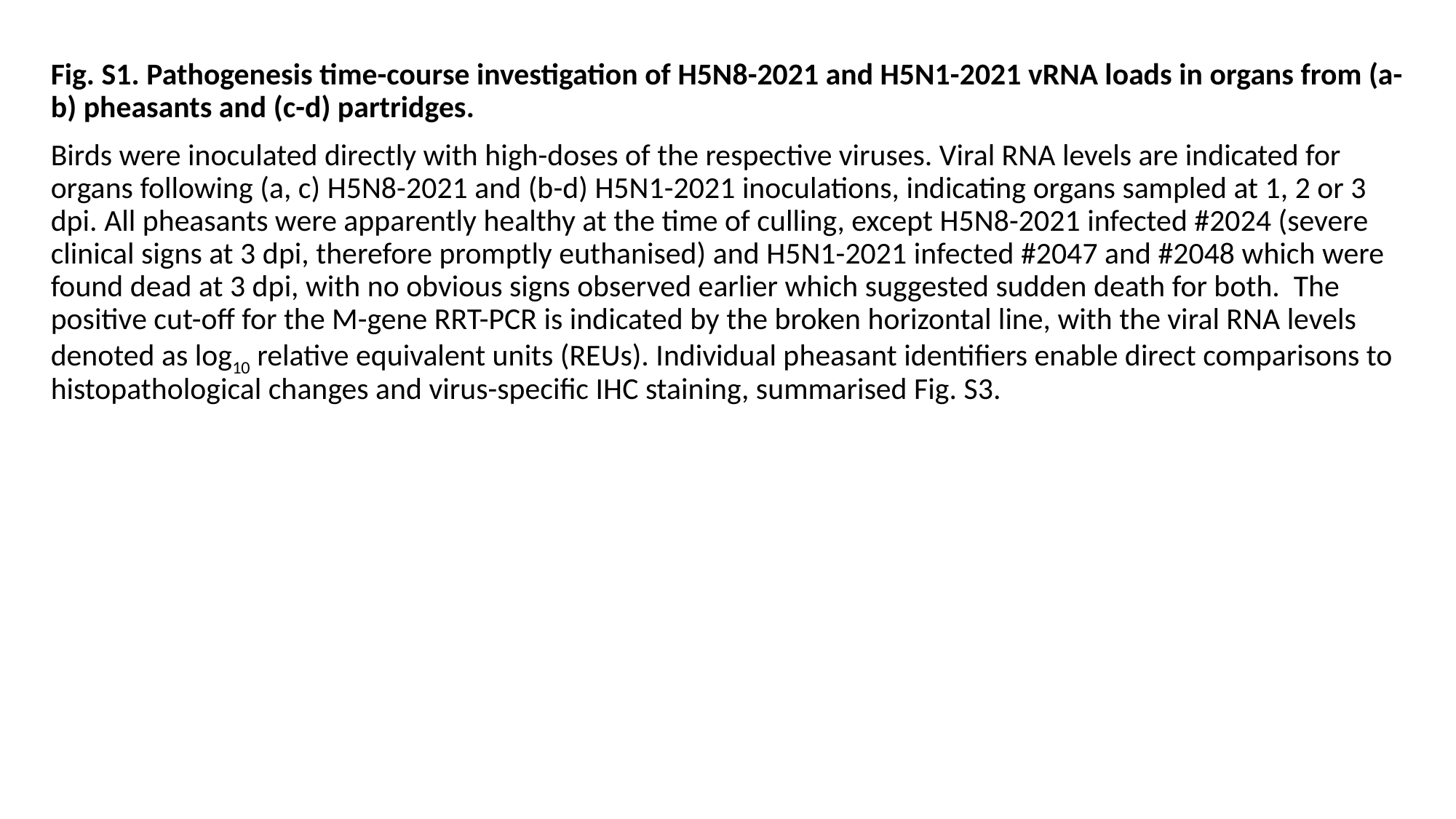

Fig. S1. Pathogenesis time-course investigation of H5N8-2021 and H5N1-2021 vRNA loads in organs from (a-b) pheasants and (c-d) partridges.
Birds were inoculated directly with high-doses of the respective viruses. Viral RNA levels are indicated for organs following (a, c) H5N8-2021 and (b-d) H5N1-2021 inoculations, indicating organs sampled at 1, 2 or 3 dpi. All pheasants were apparently healthy at the time of culling, except H5N8-2021 infected #2024 (severe clinical signs at 3 dpi, therefore promptly euthanised) and H5N1-2021 infected #2047 and #2048 which were found dead at 3 dpi, with no obvious signs observed earlier which suggested sudden death for both. The positive cut-off for the M-gene RRT-PCR is indicated by the broken horizontal line, with the viral RNA levels denoted as log_10_ relative equivalent units (REUs). Individual pheasant identifiers enable direct comparisons to histopathological changes and virus-specific IHC staining, summarised Fig. S3.

### Slide 3
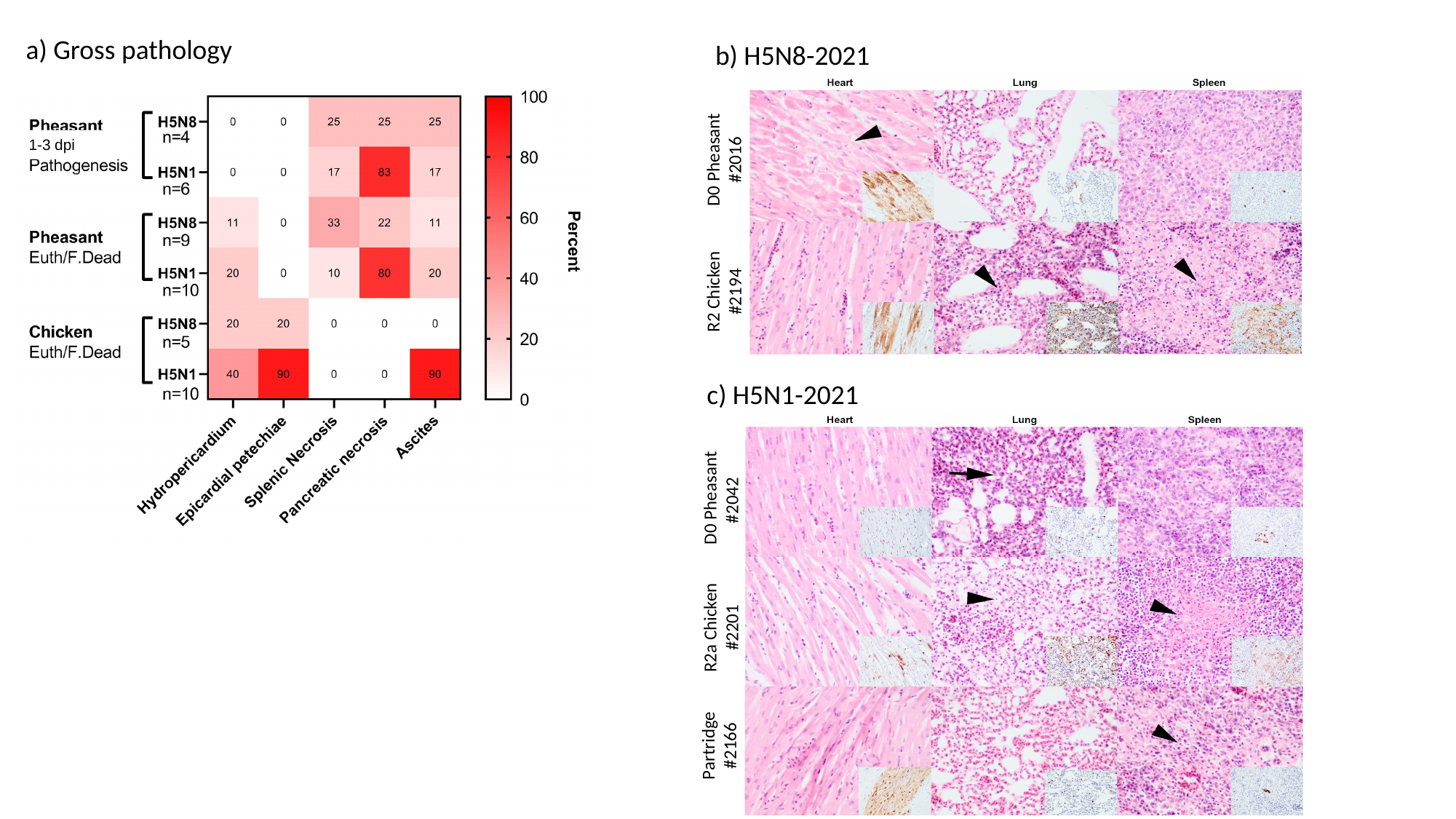

a) Gross pathology
b) H5N8-2021
D0 Pheasant
#2016
R2 Chicken
#2194
1-3 dpi
c) H5N1-2021
 D0 Pheasant
#2042
R2a Chicken
#2201
Partridge
#2166

### Slide 4
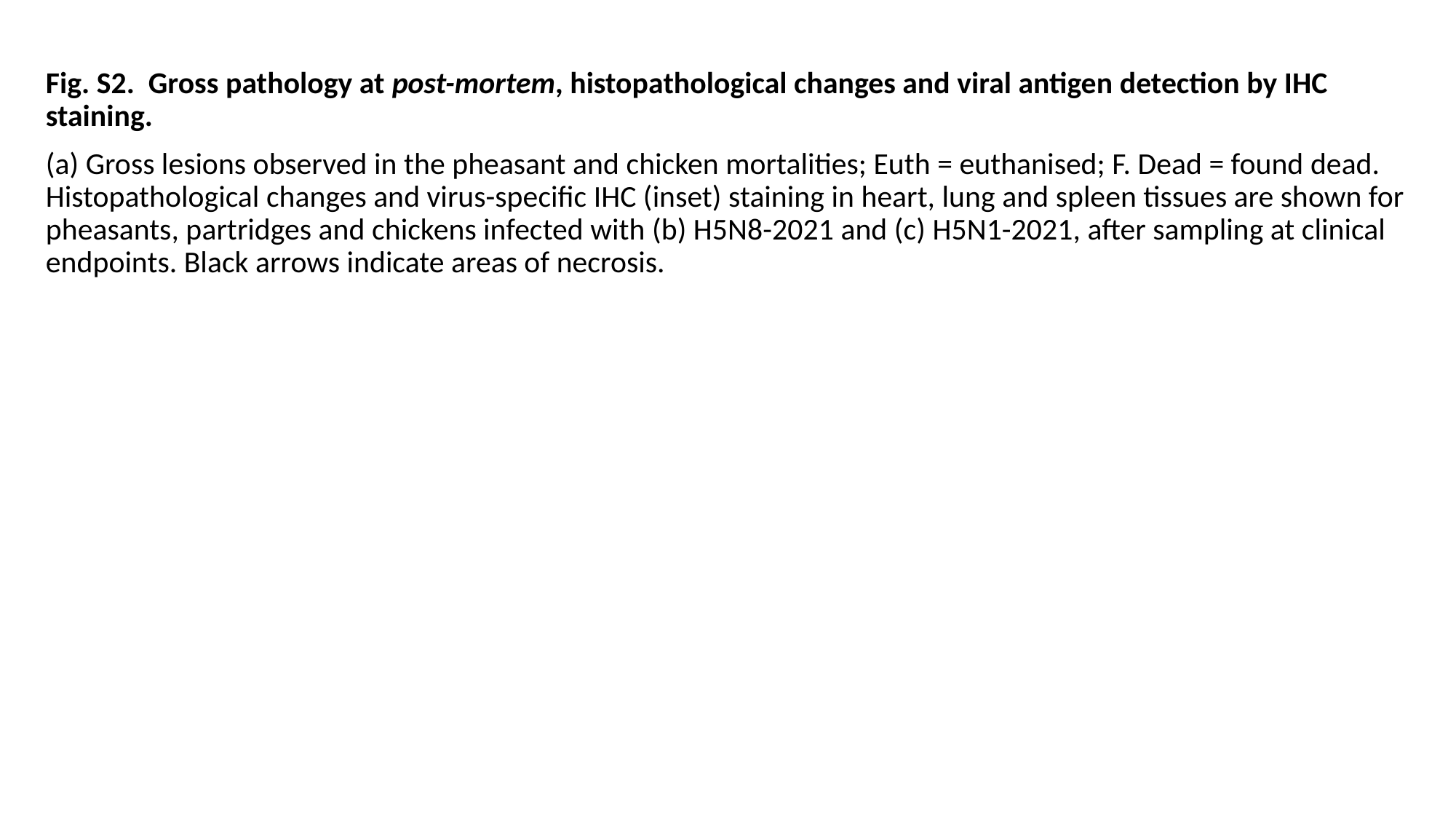

Fig. S2. Gross pathology at post-mortem, histopathological changes and viral antigen detection by IHC staining.
(a) Gross lesions observed in the pheasant and chicken mortalities; Euth = euthanised; F. Dead = found dead. Histopathological changes and virus-specific IHC (inset) staining in heart, lung and spleen tissues are shown for pheasants, partridges and chickens infected with (b) H5N8-2021 and (c) H5N1-2021, after sampling at clinical endpoints. Black arrows indicate areas of necrosis.

### Slide 5
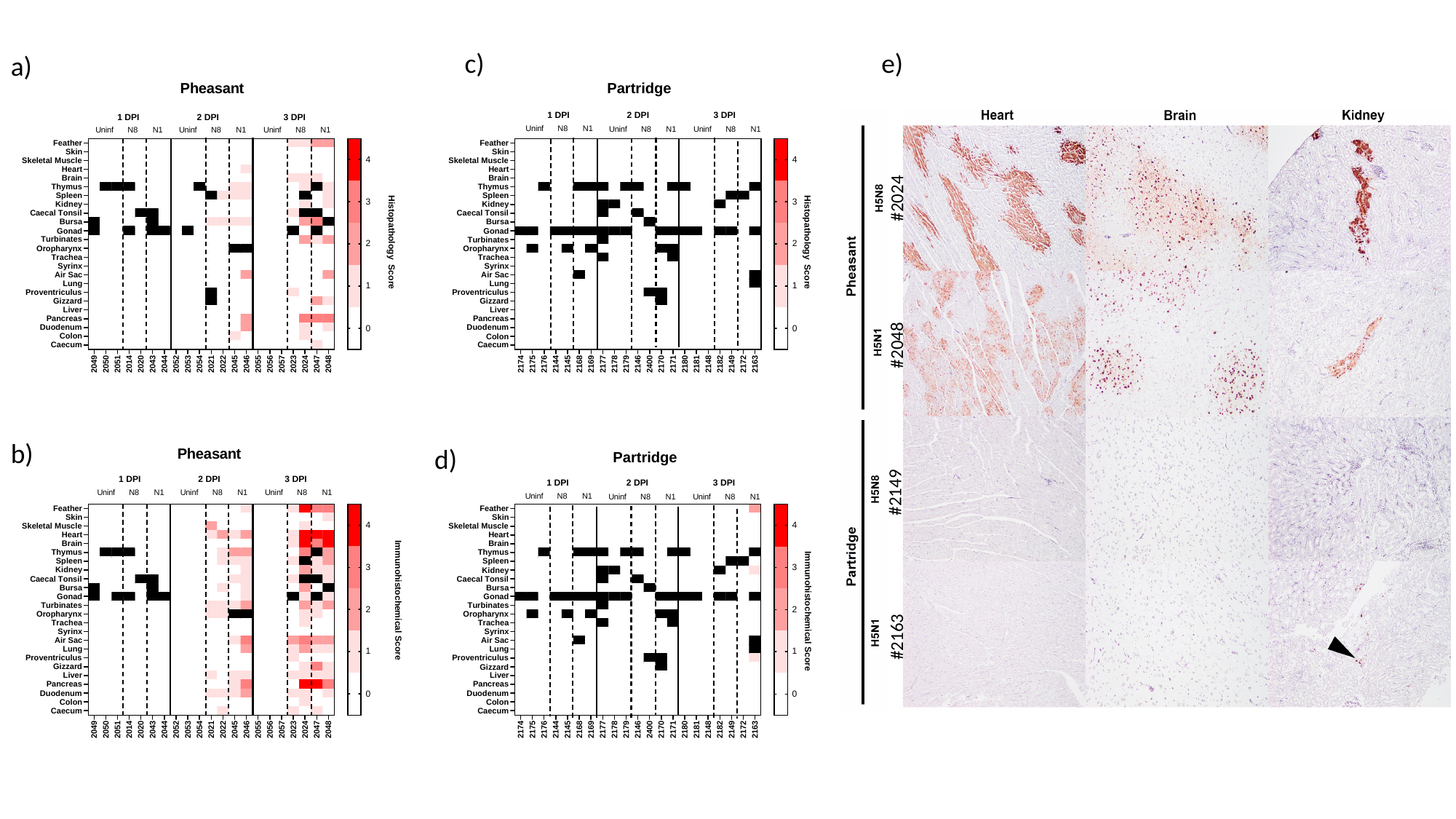

e)
c)
a)
#2024
#2048
#2149
#2163
b)
d)

### Slide 6
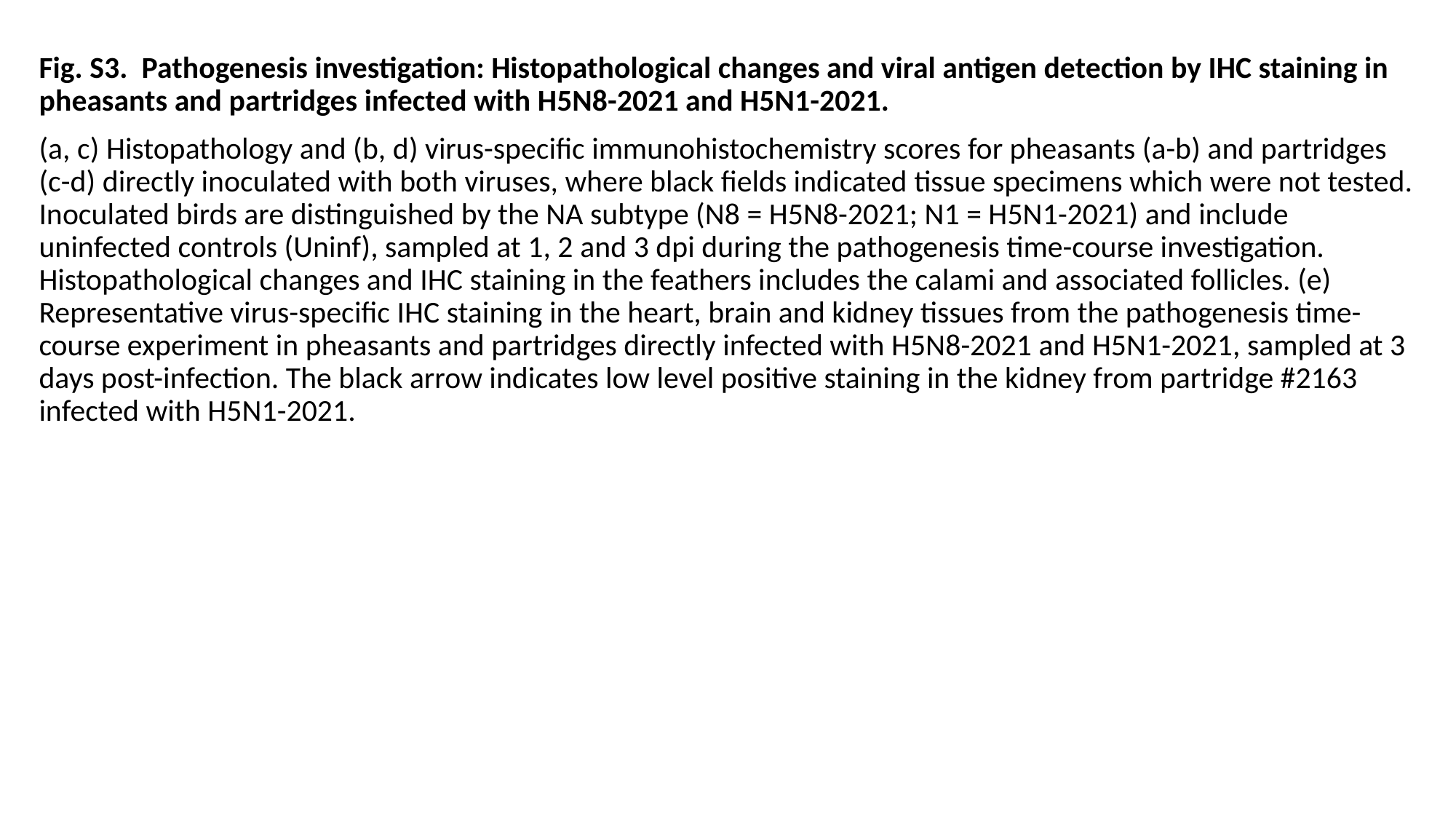

Fig. S3. Pathogenesis investigation: Histopathological changes and viral antigen detection by IHC staining in pheasants and partridges infected with H5N8-2021 and H5N1-2021.
(a, c) Histopathology and (b, d) virus-specific immunohistochemistry scores for pheasants (a-b) and partridges (c-d) directly inoculated with both viruses, where black fields indicated tissue specimens which were not tested. Inoculated birds are distinguished by the NA subtype (N8 = H5N8-2021; N1 = H5N1-2021) and include uninfected controls (Uninf), sampled at 1, 2 and 3 dpi during the pathogenesis time-course investigation. Histopathological changes and IHC staining in the feathers includes the calami and associated follicles. (e) Representative virus-specific IHC staining in the heart, brain and kidney tissues from the pathogenesis time-course experiment in pheasants and partridges directly infected with H5N8-2021 and H5N1-2021, sampled at 3 days post-infection. The black arrow indicates low level positive staining in the kidney from partridge #2163 infected with H5N1-2021.

### Slide 7
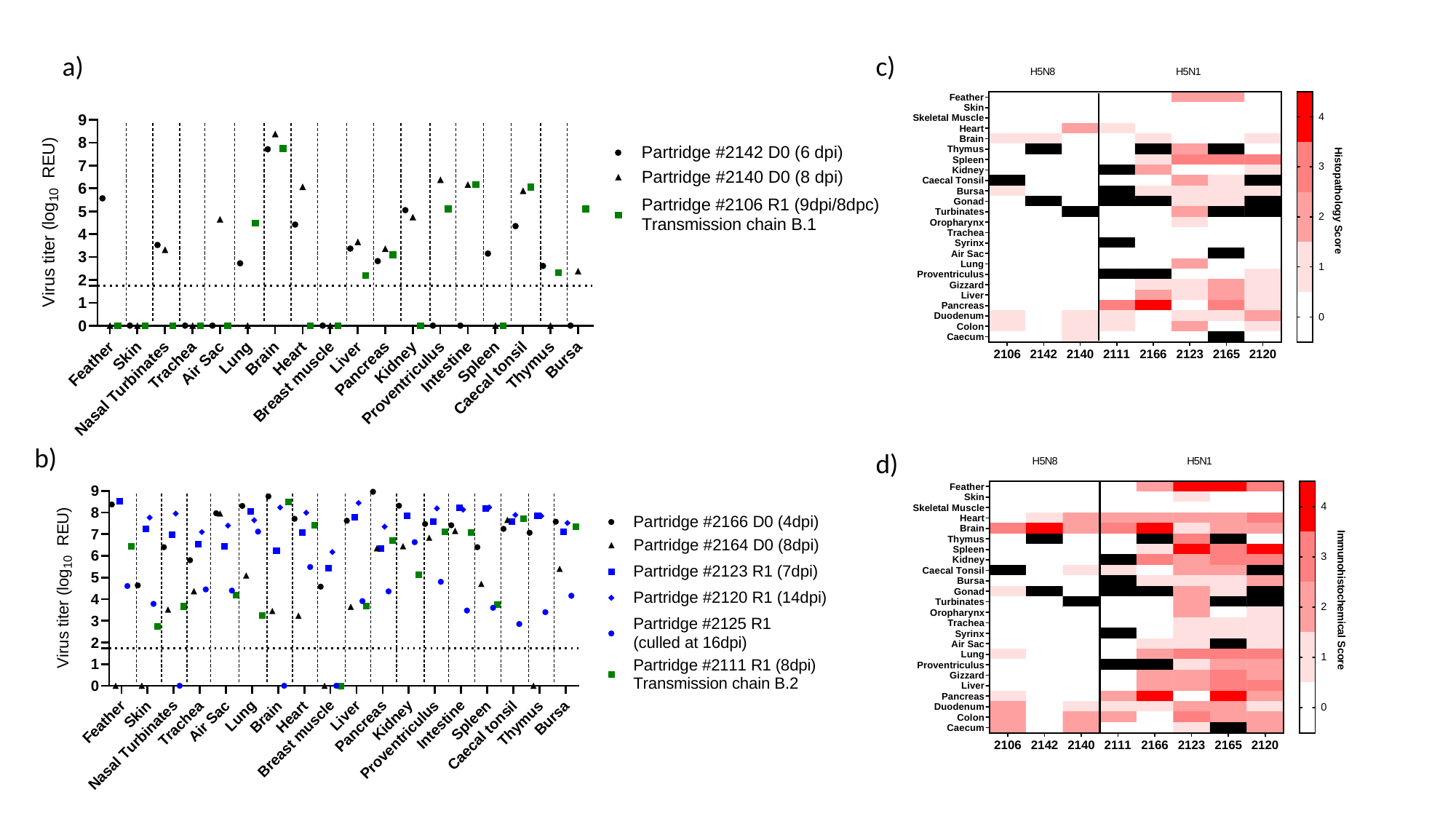

a)
c)
b)
d)

### Slide 8
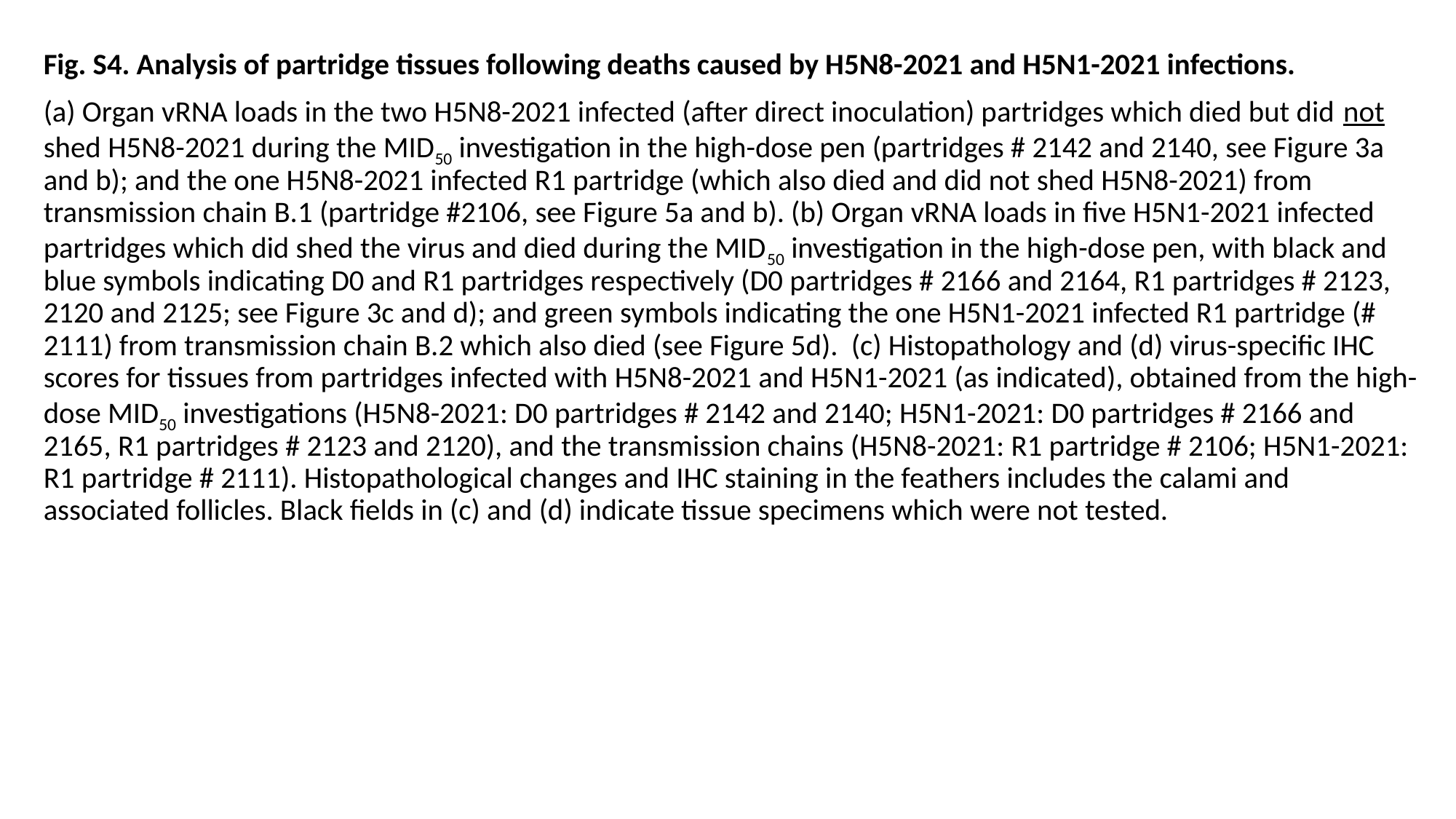

Fig. S4. Analysis of partridge tissues following deaths caused by H5N8-2021 and H5N1-2021 infections.
(a) Organ vRNA loads in the two H5N8-2021 infected (after direct inoculation) partridges which died but did not shed H5N8-2021 during the MID_50_ investigation in the high-dose pen (partridges # 2142 and 2140, see Figure 3a and b); and the one H5N8-2021 infected R1 partridge (which also died and did not shed H5N8-2021) from transmission chain B.1 (partridge #2106, see Figure 5a and b). (b) Organ vRNA loads in five H5N1-2021 infected partridges which did shed the virus and died during the MID_50_ investigation in the high-dose pen, with black and blue symbols indicating D0 and R1 partridges respectively (D0 partridges # 2166 and 2164, R1 partridges # 2123, 2120 and 2125; see Figure 3c and d); and green symbols indicating the one H5N1-2021 infected R1 partridge (# 2111) from transmission chain B.2 which also died (see Figure 5d). (c) Histopathology and (d) virus-specific IHC scores for tissues from partridges infected with H5N8-2021 and H5N1-2021 (as indicated), obtained from the high-dose MID_50_ investigations (H5N8-2021: D0 partridges # 2142 and 2140; H5N1-2021: D0 partridges # 2166 and 2165, R1 partridges # 2123 and 2120), and the transmission chains (H5N8-2021: R1 partridge # 2106; H5N1-2021: R1 partridge # 2111). Histopathological changes and IHC staining in the feathers includes the calami and associated follicles. Black fields in (c) and (d) indicate tissue specimens which were not tested.
