## Supplementary material for "Transmission dynamics and pathogenesis differ between pheasants and partridges infected with clade 2.3.4.4b H5N8 and H5N1 high-pathogenicity avian influenza viruses": Suppl. Table S1

**Table S1.** **Genetic polymorphisms which emerged relative to the (a) H5N8-2021 or (b) H5N1-2021 inocula, identified by successful WGS of progeny viruses obtained from clinical specimens at different transmission stages, from each species. Amino acid changes are emphasised by bold type and parenthesis.**

(a) H5N8-2021 mutations:

| **Experiment** | **Bird no.** | **Bird species** | **Sample** | **Infection stage** | **dpi/dpc** | **PB2** | **PB1** | **PB1-F2** | **PA** | **PA-X** | **HA** | **NP** | **NA** | **M1** | **M2** | **NS1** | **NS2** | Total  substitutions per bird |
| --- | --- | --- | --- | --- | --- | --- | --- | --- | --- | --- | --- | --- | --- | --- | --- | --- | --- | --- |
| H5N8 MID_50_ pheasants | 2011 | pheasant | buccal swab | D0 medium dose | 9 dpi | T145G **(W49G)** | - | - | - | - | - | T942C | - | - | - | - | - | 2 **(1)** |
|  | 2012 | pheasant | buccal swab | D0 medium dose | 10 dpi | - | - | - | - | - | G1536A | - | T363C | - | - | - | G237A | 3 |
| Transmission chain A.1 | 2059 | pheasant | buccal swab | D0 high dose | 4 dpi | - | - | - | - | - | - | - | - | - | - | - | - | 0 |
|  | 2063 | pheasant | buccal swab | D0 high dose | 4 dpi | - | - | - | - | - | - | - | - | - | - | - | - | 0 |
|  | 2082 | pheasant | buccal swab | R1 (ph-ph) | 6 dpi (5 dpc) | - | - | - | - | - | - | G1122A **(M374I)** | - | - | - | - | - | 1 **(1)** |
|  | 2087 | pheasant | buccal swab | R1 (ph-ph) | 6 dpi (5 dpc) | - | - | - | - | - | - | - | T228C | - | - | - | - | 1 |
|  | 2192 | chicken | buccal swab | R2  (ph-ph-ch) | 6 dpi (2 dpc) | - | - | - | - | - | - | G1122A **(M374I)** | - | - | - | - | - | 1 **(1)** |
|  | 2193 | chicken | buccal swab | R2  (ph-ph-ch) | 7 dpi (3 dpc) | - | - | - | - | - | - | - | - | - | - | - | - | 0 |
|  | 2194 | chicken | buccal swab | R2  (ph-ph-ch) | 8 dpi (4 dpc) | A693G | T1392C | - | - | - | - | T30C | - | - | - | - | - | 3 |
| Transmission chain B.1 | 2065 | pheasant | buccal swab | D0 high dose | 4 dpi | - | - | - | - | - | - | - | - | - | - | - | - | 0 |
|  | 2068 | pheasant | buccal swab | D0 high dose | 4 dpi | - | - | - | - | - | - | - | - | - | - | - | - | 0 |
|  | 2106 | partridge | brain tissue | R1 (ph-pa) | 9 dpi (8 dpc) | - | - | - | A1042G **(I348V)** | - | - | C248A **(P83H)** | - | - | - | - | - | 2 **(2)** |
| H5N8 MID_50_ partridges | 2142 | partridge | brain tissue | D0 high dose | 6 dpi | - | G1734A | - | - | - | - | - | - | - | - | - | - | 1 |
| **Total nucleotide and amino acid substitutions per segment:** | | | | | | 2 **(1)** | 2 | 0 | 1 **(1)** | 0 | 1 | 5 **(3)** | 2 | 0 | 0 | 0 | 1 |  |

(b) H5N1-2021 mutations:

| **Experiment** | **Bird no.** | **Bird species** | **Sample** | **Infection stage** | **dpi/dpc** | **PB2** | **PB1** | **PB1-F2** | **PA** | **PA-X** | **HA** | **NP** | **NA** | **M1** | **M2** | **NS1** | **NS2** | **Total substitutions per bird** |
| --- | --- | --- | --- | --- | --- | --- | --- | --- | --- | --- | --- | --- | --- | --- | --- | --- | --- | --- |
| Transmission chain A.2 | 2072 | pheasant | buccal swab | D0 high dose | 3 dpi | - | - | - | - | - | - | - | - | - | - | - | - | 0 |
|  | 2075 | pheasant | buccal swab | D0 high dose | 3 dpi | - | - | - | - | - | - | - | - | - | - | - | - | 0 |
|  | 2089 | pheasant | buccal swab | R1 (ph-ph) | 5 dpi (4 dpc) | - | - | - | - | - | - | - | - | - | - | - | - | 0 |
|  | 2386 | pheasant | buccal swab | R1 (ph-ph) | 6 dpi (5 dpc) | - | - | - | - | - | - | - | - | - | - | - | - | 0 |
|  | 2198 | chicken | buccal swab | R2a (ph-ph-ch) | 4 dpi (1 dpc) | A1425G | - | - | A6G | - | - | - | - | - | - | - | - | 2 |
|  | 2200 | chicken | buccal swab | R2a (ph-ph-ch) | 5 dpi (2 dpc) | A1425G | - | - | A6G | - | - | - | - | - | - | - | - | 2 |
|  | 2218 | chicken | buccal swab | R2b (ph-ph-ch) | 12 dpi (7 dpc) | A1425G | - | - | A6G | - | - | - | - | - | - | - | - | 2 |
|  | 2219 | chicken | buccal swab | R2b (ph-ph-ch) | 11 dpi (6 dpc) | A1425G | - | - | A6G | - | - | - | - | - | - | - | - | 2 |
|  | 2233 | chicken | buccal swab | R3 (ph-ph-ch-ch) | 12 dpi (6 dpc) | G1376A **(G459E)**, A1425G | - | - | A6G | - | - | - | - | - | - | - | - | 3 **(1)** |
|  | 2236 | chicken | buccal swab | R3 (ph-ph-ch-ch) | 11 dpi (5 dpc) | A1425G | - | - | A6G | - | - | - | - | - | - | - | - | 2 |
|  | 2238 | chicken | buccal swab | R3 (ph-ph-ch-ch) | 11 dpi (5 dpc) | A1425G | - | - | A6G | - | - | - | - | - | - | - | - | 2 |
| Transmission chain B.2 | 2078 | pheasant | buccal swab | D0 high dose | 3 dpi | - | - | - | - | - |  |  | - | - | - | - | - | 0 |
|  | 2080 | pheasant | buccal swab | D0 high dose | 3 dpi | - | - | - | - | - |  |  | - | - | - | - | - | 0 |
|  | 2111 | partridge | buccal swab | R1 (ph-pa) | 8 dpi (7 dpc) | - | - | - | - | - | A1302G | G570A | - | - | - | - | - | 2 |
|  | 2111 | partridge | brain tissue | R1 (ph-pa) | 8 dpi (7 dpc) | - | - | - | - | - | - | G570A | - | - | - | - | - | 1 |
| H5N1 MID_50_ partridges | 2162 | partridge | buccal swab | D0 high dose | 15 dpi | G546A | - | - | - | - | - | T528C, A740G **(N247S)**, G856T **(V286F)**, G1239A | - | - | - | - | - | 5 **(2)** |
|  | 2167 | partridge | buccal swab | D0 high dose | 13 dpi | G546A | - | - | - | - | - | T528C, A740G **(N247S)**, G856T **(V286F)**, G1239A | - | - | - | - | - | 5 **(2)** |
| Partridge-Partridge transmission | 2122 | partridge | buccal swab | R1 (pa-pa) | 15 dpi  (14 dpc) | G546A | - | - | - | - | - | T528C, A740G **(N247S)**, G856T **(V286F)**, G1239A | - | - | - | - | - | 5 **(2)** |
|  | 2094 | partridge | buccal swab | R1 (pa-pa) | 16 dpi  (15 dpc) | G546A | - | - | - | - | - | T528C, A740G **(N247S)**, G856T **(V286F)**, G1239A | - | - | - | - | - | 5 **(2)** |
|  | 2125 | partridge | lung tissue | R1 (pa-pa) | 16 dpi  (15 dpc) | G546A | - | - | - | - | - | T528C, A740G **(N247S)**, G856T **(V286F)**, G1239A | - | - | - | - | - | 5 **(2)** |
| **Total nucleotide and amino acid substitutions per segment:** | | | | | | 3 **(1)** | 0 | 0 | 1 | 0 | 1 | 5 **(2)** | 0 | 0 | 0 | 0 | 0 |  |
